# Enhanced cardiac mitochondrial biogenesis by nitro-oleic acid remedies diastolic dysfunction in a mouse model of heart failure with preserved ejection fraction

**DOI:** 10.1101/2024.02.20.581137

**Authors:** Marion Müller, Torben Schubert, Cornelius Welke, Tina Johanna Schulz, Thomas Patschkowski, Tibor Maske, Luisa Andrea Lengenfelder, Lucia Landwehrjohann, Elfi Donhauser, Elisa Theres Vogt, Bernd Stratmann, Jurek Hense, Simon Lüdtke, Martina Düfer, Elen Tolstik, Johann Dierks, Felix-Levin Hormann, Sven Heiles, Kristina Lorenz, Jan-Christian Reil, Francisco Jose Schopfer, Bruce A. Freeman, Volker Rudolph, Uwe Schlomann, Anna Klinke

**Affiliations:** Clinic for General and Interventional Cardiology/ Angiology, Herz- und Diabeteszentrum NRW, University Hospital of the Ruhr-Universität Bochum, Bad Oeynhausen, Germany; Agnes Wittenborg Institute for Translational Cardiovascular Research (AWIHK), Herz- und Diabeteszentrum NRW, University Hospital of the Ruhr-Universität Bochum, Bad Oeynhausen, Germany; Technology Platform Genomics, Center for Biotechnology (CeBiTec), Bielefeld University, Bielefeld, Germany; Diabetescenter, Herz- und Diabeteszentrum NRW, University Hospital of the Ruhr-Universität Bochum, Bad Oeynhausen, Germany; Institute of Pharmaceutical and Medicinal Chemistry, University of Münster, Münster, Germany; Leibniz-Institut für Analytische Wissenschaften-ISAS e.V., Dortmund, Germany; Faculty of Chemistry, University of Duisburg-Essen, 45141 Essen, Germany; Institute of Pharmacology and Toxicology, University of Würzburg, Würzburg, Germany; Department of Pharmacology and Chemical Biology, University of Pittsburgh, Pittsburgh, PA, USA

## Abstract

Prevalence of heart failure with preserved ejection fraction (HFpEF) is increasing, while treatment options are inadequate. Hypertension and obesity-related metabolic dysfunctions contribute to HFpEF progression. Nitro-oleic acid (NO_2_-OA) impacts metabolic processes by improving glucose tolerance and adipocyte function. In this study, 4 week treatment with NO_2_-OA ameliorated diastolic dysfunction in a HFpEF mouse model induced by high-fat diet and inhibition of the endothelial nitric oxide synthase. A proteomic analysis of left ventricular tissue revealed, that one third of the identified proteins, mostly mitochondrial proteins, were upregulated in hearts of NO_2_-OA-treated HFpEF mice compared to controls and vehicle-treated HFpEF mice, which was confirmed by immunoblot. Activation of the 5’-adenosine-monophosphate-activated-protein-kinase (AMPK) signaling pathway mediated an enhancement of mitochondrial biogenesis in hearts of NO_2_-OA-treated HFpEF mice. In cardiomyocytes under metabolic stress, NO_2_-OA increased mitochondrial protein level accompanied by enhanced oxidative phosphorylation. In conclusion, targeting mitochondrial integrity in HFpEF leads to improved diastolic function.

## Introduction

Heart failure with preserved ejection fraction (HFpEF) is a disorder with high morbidity and mortality and an increasing prevalence in the aging populations of Western countries^1^. Given the heterogeneity and multifactorial nature of this syndrome, the definition of distinct phenogroups has been established recently to better stratify HFpEF patients^2–4^. The group of patients with cardiometabolic HFpEF exhibits obesity and metabolic syndrome. Systemic vascular inflammation and hypertension are further common phenomena of this phenogroup. Sodium-glucose cotransporter 2 (SGLT2) inhibitors reduce the risk of cardiovascular death in HFpEF^5,6^, while the mechanisms of action remain elusive. Apart from that, pharmacological options that directly target cardiac malfunction are not available, given that even in a particular phenogroup the pathogenesis of diastolic dysfunction is multifactorial and incompletely understood. One pathophysiological hallmark is metabolic remodeling of the myocardium, insulin resistance and altered lipid handling. This is driven by elevated lipid levels, which promotes a decrease in glucose uptake and consequent hyperinsulinemia^7^. This, together with multiple other risk factors such as physical inactivity, stimulates compensatory cardiomyocyte hypertrophy and causes microvascular dysfunction, resulting in reduced oxygen supply to the cardiomyocyte. However, oxygen is required during aerobic energy generation. Fatty acid oxidation (FAO) consumes more oxygen than glucose oxidation (GO) or glycolysis. At the same time, FAO is the most effective way to generate adenosine triphosphate (ATP) per molecule substrate, with the released energy being proportional to acyl chain length. The metabolic dysregulation of HFpEF impairs metabolic flexibility, so that mitochondrial metabolism cannot adapt to the variable cardiomyocyte demand for energy and oxygen. Given that insulin resistance shifts the cardiomyocyte substrate metabolism further towards FAO, the role of FAO in cardiometabolic HFpEF remains controversial^8,9^. In fact, lipotoxicity has been linked to the HFpEF pathophysiology. It is characterized by cardiomyocyte accumulation of lipid intermediates such as ceramide and diacylglycerol (DAG) due to the increased uptake of fatty acid (FA) and or a decreased rate of FAO^8,10^. Importantly, alterations in the lipid composition of cell and organelle membranes can disturb cellular integrity and also lead to mitochondrial dysfunction^11,12^. Critical mediators of metabolic regulation in cardiomyocytes include AMP-activated protein kinase (AMPK) and the deacetylase sirtuin 1 (SIRT1), which promote catabolic pathways, in particular FAO, in part via activation of mitochondrial biogenesis and the peroxisome-proliferator activated receptor (PPAR)-α.

Nitro-fatty acids are electrophilic molecules formed by reactions of nitrogen oxides, particularly nitrogen dioxide, with FA containing conjugated double bonds. Both endogenous and synthetic electrophilic nitroalkene derivatives of FA react with functionally significant cysteine moieties in enzymes and transcription factors to modulate protein function and induce salutary signaling responses. Free endogenous plasma levels under basal conditions are ∼1-3 nM, and can increase during inflammation and through dietary interventions to reach levels of 7-10 nM^13^. Similar levels are achieved in animal models such as herein and in clinical studies when orally dosing human subjects with 150 mg NO_2_-OA^14,15^. Notably, the free acid levels reflected by these plasma measurements do not include the esterified pool of nitro-fatty acids, which is ∼20-30 fold greater than the free acid levels in plasma^16^. Experimental studies have extensively explored the effects of the synthetic isomers (*E*)-9- and 10-nitrooctadec-9-enoic acid (nitro-oleic acid, NO_2_-OA) and (9*E*,12*E*)-9-nitrooctadeca-9,12-dienoic acid (nitro-linoleic acid, NO_2_-LA) or the predominant endogenous conjugated (9*E*,11*E*)-linoleic acids (NO_2_-CLA)^17–19^. An ongoing phase 2 clinical trial evaluates the anti-inflammatory actions of (*E*)-10-nitrooctadec-9-enoic acid on airway dysfunction in obese asthmatics (NCT03762395). Small molecule nitroalkenes exert anti-inflammatory and antioxidant properties in part via activation of the nuclear factor erythroid 2-related factor 2 (Nrf2) and inhibition of the nuclear factor ‘kappa-light-chain-enhancer’ of activated B-cells (NF-κB)^20^. Interestingly, they also improve energy metabolism, not only by limiting the pro-inflammatory milieu in liver and adipose tissue^21,22^, but also by ligand-mediated activation of PPARs^23^. In leptin-deficient mice, obesity, glucose intolerance, and hyperglycemia were reduced by subcutaneous NO_2_-OA administration^24^. In a high fat diet (HFD) mouse model, animals were protected from right ventricular dysfunction by NO_2_-OA via reduction of pulmonary arterial remodeling^21^.

The ‘two-hit’ HFpEF mouse model of combined HFD with *N*_ω_-nitro-L-arginine methyl ester hydrochloride (L-NAME) administration to inhibit the endothelial nitric oxide synthase (eNOS), established by Schiattarella and colleagues^25^, unites the two major HFpEF risk factors - hypertension and metabolic syndrome. We employed this 15-week HFD+L-NAME model to study LV function responses to NO_2_-OA administered in the last 4 weeks of treatment.

## Methods

### Animals

Male C57BL/6N mice were purchased from Charles River (Sulzfeld, Germany). Mice were housed for at least 1 week to acclimatize to laboratory conditions before starting any experimental procedure. They were kept in a 12:12 hours inverse light cycle and had unrestricted access to water and food.

All animal studies were approved by the local Animal Care and Use Committees (Ministry for Environment, Agriculture, Conservation and Consumer Protection of the State of North Rhine-Westphalia: State Agency for Nature, Environment and Consumer Protection (LANUV), NRW, Germany) and followed guidelines from Directive 2010/63/EU of the European Parliament on the protection of animals used for scientific purposes.

### Animal protocols

Cardiometabolic syndrome was induced in mice by HFD combined with the established eNOS inhibitor L-NAME as described by Schiattarella *et al.*^25^. 4-week-old mice were fed a HFD (60% fat, 20% protein, 20% carbohydrates; E15742-34; ssniff, Germany) or standard rodent diet (chow group; V1534-703; ssniff, Germany). L-NAME (0.5 g/L; #N5751; Sigma Aldrich) was supplied in the drinking water. The HFD+L-NAME and chow groups were fed for 11 weeks and body weight was determined every week. After 11 weeks, the HFD+L-NAME group was divided into 3 groups with equal range of body weight (mean ± SD: HFD+L-NAME vehicle: 42.4 ± 5.9 g; HFD+L-NAME OA: 42.5 ± 4.0 g; HFD+L-NAME NO_2_-OA: 43.6 ± 3.6 g) and treated over 4 weeks, while HFD+L-NAME was continued. Nitro-oleic acid (50:50 mix of (*E*)-9- and 10-nitrooctadec-9-enoic acid; NO_2_-OA, was synthesized *in-house* as previously^26^, oleic acid (OA) or vehicle (polyethylene glycol:ethanol 90:10) were administered with ALZET mini-osmotic pumps (NO_2_-OA and OA 8.17 mg/kg bw/day, Model 2002, ALZET, Cupertino, CA, USA). Cardiac ultrasound was performed before (11 weeks, baseline) and after 2 (13 weeks) and 4 weeks (15 weeks) of treatment as described below. After final investigations, organs were harvested and snap-frozen for RNA and protein analysis and fixed in formalin.

### Echocardiography

Cardiac ultrasound was carried out with a Vevo 3100 Imaging System (FUJIFILM Visualsonics, Inc., Toronto, ON, Canada) using a MX550D transducer (25-55 MHz, FUJIFILM Visualsonics, Inc., Toronto, ON, Canada). Mice were anaesthetized with isoflurane inhalation (1.5-2%) and placed in supine position on a heating pad. An electrocardiogram was obtained with integrated electrodes. Body temperature was monitored using a rectal probe (T = 36.5-37.5 °C). Respiration rate (RR = 80-120) was controlled by adjusting the depth of anesthesia. A standardized workflow was followed as stated in the position paper of the Working Group on Myocardial Function of the European Society of Cardiology^27^. In brief, 2D recordings of brightness- (B-) and motion- (M-) mode of parasternal long-axis (PSLAX) and parasternal short-axis (PSAX), respectively, were acquired for analysis of LV ejection fraction (LVEF) and posterior wall thickness in diastole (LVPW, d). Diastolic function was obtained in apical four-chamber view (4APIX) with pulsed-wave and tissue Doppler imaging at the level of the mitral valve. Peak Doppler blood inflow velocity across the mitral valve during early diastole (E wave) and during late diastole (A wave) and peak tissue Doppler of myocardial relaxation velocity at the mitral valve annulus during early diastole (e’ wave) were assessed. All parameters were determined three times. The average is presented for each animal.

### Strain analysis

The strain analysis was carried out using the Vevo strain analysis module of the Vevo Lab analysis software (Version 3.2.0; FUJIFILM Visualsonics, Inc., Toronto, ON, Canada). Strain determines the deformation of the myocardial wall and thereby reflects cardiac function. The analysis uses two-dimensional speckle tracking to calculate the distance a speckle moves between two consecutive frames, which is indicated as displacement. The velocity is the displacement per unit time in three planes. The longitudinal strain is a dimensionless parameter and shows the tangential movement based on the traced border normalized towards a baseline. The software uses the Lagrangian Strain algorithm, which is based on the following formula: S(t) = (L(t) – L(0)) / L(0); S(t), Strain value at a specific time point t; L(t), length at a specific time point t; L(0), length at the start point 0. The analysis was performed by using three cardiac cycles of recordings of parasternal long-axis (PSLAX) view.

### Glucose-tolerance test

Glucose-tolerance tests were performed before final harvest at 15 weeks of HFD+L-NAME. After 6 h fastening glucose solution (2 g/kg in saline) was injected intraperitoneally. Tail blood glucose levels (mg/dL) were measured with a glucometer (B Braun Melsungen, Melsungen, Germany) before (0 min) and at 15 min and 90 min after glucose administration. The area under the curve (AUC) was calculated by using the series of determined measurements over time.

### Liquid chromatography-mass spectrometry (LC-MS)

The LV tissue was ground in liquid nitrogen to a fine powder. After incubation for 10 minutes in 4% SDS Tris-Cl buffer (pH 7.6) and centrifugation (16,000 g for 15 minutes), the supernatants were diluted with 4 volumes of ice-cold acetone to precipitate the proteins (overnight at −20 °C). The proteins were redissolved in 8 M Urea/50 mM TEAB buffer containing protease- and phosphatase inhibitors (EDTA free cOmplete^TM^, PhosSTOP^TM^ Roche) and treated 30 minutes at 37 °C with 50 units/100 µg tissue benzonase (Sigma Aldrich) to degrade chromatin. A BCA Assay (Pierce) was used to determine the protein concentration. After reduction of each 100 µg Protein (5 mM DTT for 1h at 25 °C) and alkylation (40 mM iodacetamide for 30 minutes at 25 °C in the dark), Lys-C (Promega) was added in an enzyme:substrate ratio of 1:75 for 4 hours at 25 °C. The samples were diluted with 50 mM TEAB buffer to a final Urea concentration of 2 M before adding trypsin (Promega) in an enzyme:substrate ratio of 1:75 and incubating at 25 °C for 16 hours. The enzymatic digestion was stopped by adding formic acid to a final concentration of 1%. The stop and Go extraction (Stage) was used to prepare the protein digestions for mass spectrometric analysis (Rappsilber et al., 2003). Two small pieces of Empore^TM^ SDB-RPS extraction disks (Supelco, 66886-U) were placed with a 16 gauge blunt-ended syringe needle in 200 µl pipette tips. StageTips were conditioned by 60 µl of methanol and centrifugation at 450 g for 75 seconds, followed by centrifugation with 60 µl buffer B (0.1% formic acid / 80% acetonitrile) and two times with 60 µl buffer A (0.1% formic acid) for equilibration. The samples were loaded and passed through the StageTips by centrifugation at 450 g for 75 seconds, followed by washing steps with 60 µl buffer A and two times with 60 µl buffer B. After air drying, the samples were eluted two times with 60 µl 60:35:5% acetonitrile:water:ammonium hydroxide, vacuum dried, and redissolved in 50% acetonitrile / 0.1% TFA in a concentration of 1 µg peptide/µl.

The peptides were analysed using a nanoLC (Ultimate 3000, Thermo Fisher Scientific, Germany) coupled to an ESI-Orbitrap MS/MS (QExactive Plus, Thermo Fisher Scientific, Germany). The gradient length of the Acclaim™ PepMap™ C18 2UM 75UMx500MM (Thermo Fisher Scientific, Dreieich, Germany) analytical column was adjusted to 187 min from 4 to 50% of 80% ACN and 0.08% FA at a flow rate of 300 nl/min. ESI-Orbitrap mass spectrometry measurements were carried out in a data-dependent top-10 acquisition mode. All samples were measured in full MS mode using a resolution of 70,000 (AGC target of 3e6 and 64 ms maximum IT). For the dd-MS2 a resolution of 17,500, AGC target of 2e5 and a maximum IT of 200 ms was used. The Proteome Discoverer Software (Thermo Fisher Scientific) and Perseus (MPI Biochemistry) were used to analyse the spectra and statistical analysis.

### Cardiomyocyte isolation

Murine cardiomyocytes were isolated using a modified Langendorff apparatus as described^28^. The perfusion buffer, enzymes and enzyme concentration (100 µg/mL Liberase TM (Roche)) and stop buffers were adapted to achieve a maximum yield of rod-shaped cardiomyocytes (details given in Suppl. Material).

The aorta was canulated and the heart flushed retrograde with perfusion buffer for 3 minutes and then with digestion buffer for 3-10 minutes, depending on the drip speed. The digestion was assumed completed when the drip speed reached 3-5 drops per second. The heart was removed from the cannula, poured over with Stop Buffer, and the atria and vessels were removed and sliced into small pieces. The suspension was filtrated through a 200 µm mesh and cells were allowed to settle by gravity. Afterwards, cells were purified with solutions of increasing calcium concentrations (75, 225, 600, 1500 µM).

### Cardiomyocyte culture

The primary mouse cardiomyocytes were cultured on laminin (Merck, L2020) surface-coated four well Permanox slides (Thermo Fisher Sci, 177437) in culture medium (M199 Hanks (Gibco 12350-039), 5 mM creatine, 2 mM carnitine, 5 mM taurine, SITE+3 (Sigma S5295), 25 mM blebbistatin (Toronto Research Chemicals B592500)). To simulate a diabetic milieu^29^, the glucose concentration was increased to 10 mM, and 10 nM endothelin-1 and 1 µM hydrocortisone were added together with or without 1 µM NO_2_-OA. Methanol was used as vehicle. Culture was continued at 37 °C for 48 hrs.

### Immunoblotting

Proteins from LV tissue were isolated as described above for LC-MS. For protein isolation from isolated cardiomyocytes, a two-step discontinuous Percoll (17089101, cytiva) gradient (high density 1.086 g/ml, low density 1.060 g/ml) was used to separate the pure fraction of intact cardiomyocytes. After 45 minutes of centrifugation at 1,800 x g, the band with intact cardiomyocytes between the two layers was collected and washed three times with 4 ml PBS. The cell pellet was lysed in 50 µl RIPA buffer (150 mM NaCl, 5 mM EDTA, Tris-Cl pH 8.0, 1% NP-40, 0.5% sodium deoxycholate, 0.1% SDS) supplemented with protease inhibitor (EDTA free cOmpleteTM, Roche) as well as phosphatase inhibitor (PhosSTOPTM, Roche) for 15 minutes on ice, two times shock frozen in liquid nitrogen and afterwards centrifuged at 20,000 x g at 4 °C. Protein concentrations were determined using Pierce BCA Protein Assay Kit (23225; Thermo Fisher Scientific). Equal amounts of protein from each sample were separated on SDS– polyacrylamide gels and transferred to PVDF membrane. Membranes were stained with primary antibodies against: cytochrome-C-oxidase subunit IV (4844, Cell Signaling, Danvers, MA, USA), sirtuin 3 (5490, Cell Signaling, Danvers, MA, USA), sirtuin 1 (9475, Cell Signaling, Danvers, MA, USA) phosphorylation at threonine 172 of AMP-activated protein kinase (2531, Cell Signaling, Danvers, MA, USA), AMP-activated protein kinase (2532, Cell Signaling, Danvers, MA, USA), phosphorylation at serine 293 of pyruvate dehydrogenase (31866, Cell Signaling, Danvers, MA, USA), translocase of outer mitochondrial membrane 70 (14528-1-AP, Proteintech, Rosemont, Il, USA), pyruvate dehydrogenase kinase 4 (ab214938, abcam, Cambridge, UK) and 3-hydroxybutyrate dehydrogenase 1 (15417-1-AP, Thermo Fisher Sci, Waltham, MA, USA). HRP-conjugated secondary antibodies (NA9340, VWR International GmbH, Darmstadt, Germany) and chemiluminescent substrate (WesternBright Chemilumineszenz Substrat Quantum, Biozym, Oldendorf, Germany) was used for detection. Images were acquired using INTAS ECL CHEMOSTAR (INTAS, Göttingen, Germany), and densitometry analysis was evaluated using LabImage1D (Kapelan Bio-Imaging GmbH, Leipzig, Germany) with rolling ball background reduction. Samples were at least analysed in duplicate and values were normalized to total protein assessed by fluorescence staining (SPL Red, PR926, NH DyeAGNOSTICS, Halle (Saale), Germany). Data are shown relative to the mean of the chow group.

### Histology

Formalin-fixed, paraffin-embedded LV cross-sections of 5 µm were stained with picrosirius red (Polyscience Inc, Warrington, PA, USA) following standard protocols. Images were acquired with a BZ-X 810 microscope (Keyence, Osaka, Japan). Extent of interstitial fibrosis was graded by a blinded person in at least 6 sections of two different regions of the LV and defined as none, mild, moderate or pronounced fibrosis.

### Insulin analysis

Islets were isolated by collagenase digestion. After lysis in acidic ethanol, insulin was determined in batches of 5 islets per sample by radioimmunoassay with rat insulin as standard.

### Immunohistochemistry

For analysis of cardiomyocyte hypertrophy, paraffin-embedded 5 µm LV cross-sections were incubated with Alexa Fluor 594-conjugated wheat-germ agglutinin (WGA, Thermo Fisher Scientific, Waltham, MA, USA). Images were acquired with a BZ-X 810 microscope (Keyence, Osaka, Japan) and cardiomyocyte cross-sectional area from WGA staining was analysed using a JavaCyte macro as described previously^30^. For islet histology, paraffin-embedded 5 µm slices were stained for insulin (ab195956, 1:1500, Alexa Fluor 555-conjugated secondary antibody: ab150186, 1:500, Abcam) and glucagon (ab92517, 1:2000, Alexa Fluor 488-conjugated secondary antibody ab150077, 1:1000, Abcam). Nuclei were visualized by DAPI (Fluoroshield, Sigma Aldrich). Confocal images (10 sections, 0.35 µm distance each) were acquired by a digital inverted microscope (iMIC, FEI Germany) and evaluated by Fiji (ImageJ). The mitochondrial protein translocase of outer membrane 70 (TOM70) and the sarcomere protein Troponin T (TnT) were stained in LV cryosections of 10 µm using a rabbit antibody to TOM70 (14528-1-AP, 1:200, Proteintech; Alexa Fluor 488-conjugated secondary antibody: A-11034, 1:500, Invitrogen) and a mouse Alexa Fluor 647-conjugated antibody to TnT (#565744; BD Pharmingen, 1:100). Nuclei were stained with DAPI (1:1000, ab228549, Abcam). Confocal microscopy images were acquired using a TCS SP8 system (Leica Microsystems, Wetzlar, Germany).

### Quantitative real-time PCR

Total mRNA was isolated from murine frozen tissues using the miRNeasy Micro Kit (Qiagen, Hilden, Germany) following the manufactureŕs standard protocol. Reverse transcription was performed for 30 min at 42 °C using dNTP Mix (10 mM each, VWR, Radnor, PA, USA) and SuperScript II Reverse Transcriptase (Thermo Fisher Scientific, Waltham, MA, USA). qPCR was carried out on StepOnePlus (Applied Biosystems, Foster City, CA, USA) using Power Sybr^®^ Green qPCR Master Mix (4367659, Applied Biosystems, Foster City, CA, USA) and specific primers: *Sirt1-F*: 5’-GTGTCATAGGCTAGGTGGTGA-3’; *Sirt1-R*: 5’-TCCTTTTGTGGGCGTGGAGG-3’; *Nmnat1-F*: 5’-GTGCCCAACTTGTGGAAGAT-3’; *Nmnat1-R*: 5’-CAGCACATCGGACTCGTAGA-3’; *Rpl32-F*: 5’-CGGAAACCCAGAGGCATTGA-3’ *Rpl32-R*: 5’-GGACCAGGAACTTGCGGAAG-3’. Tomm70 mRNA expression was measured with TaqMan real-time PCR assay on StepOnePlus (Applied Biosystems, Foster City, CA, USA) using HotStarTaq DNA Polymerase (Qiagen, Hilden, Germany) according to the manufacturer’s instructions. The primer sequences are: *Tomm70-F*: 5’-AAGCGCAACAGTGAACGGAA-3’; *Tomm70-R*: 5’-CTGTCCAGAGAGCTCATTTCCA-3’; *Tomm70-P*: 5’-TGGCCACCACGACGGTTCTG-3’; *Rpl32-F*: 5’-GCTGATGTGCAACAAATCTTA-3’; *Rpl32-R: 5’-* TCGGTTCTTAGAGGACACA-3’; *Rpl32-P*: 5’-TGTGAGCAATCTCAGCAC-3’. ΔCT was calculated related to the expression of the ribosomal protein L32 (*Rpl32*). 2^-ΔΔCT^ was calculated reflecting expressions relative to the mean of the chow group.

### Mitochondrial copy number

Mitochondrial copy numbers were quantified using DNA by TaqMan real-time PCR assays. Total DNA was isolated from murine frozen tissues using the Pure Link Genomic DNA Kit (Life Technologies, Carlsbad, CA, USA) following the manufactureŕs standard protocol. RT-PCR was carried out on StepOnePlus (Applied Biosystems, Foster City, CA, USA) using HotStarTaq DNA Polymerase (Qiagen, Hilden, Germany) according to the manufacturer’s instructions. Specific primer sequences and probes targeting mitochondrial DNA (mtDNA) or genomic DNA (gDNA) were used: mtDNA-F: 5’-TCGCAGCTACAGGAAAATCA-3’; mtDNA-R: 5’-TGAAACTGGTGTAGGGCCTT-3’; mtDNA-P: 5’-GCACAATTTGGCCTCCACCCA-3’; gDNA-F: 5’-CAGAGACAGCAAACATCAGA-3’; gDNA-R: 5’-CAGGGTGATGGAGAAGGA-3’; gDNA-P: 5’-CCCGACCCACGCCAGCAT-3’. 2^-ΔΔCT^ was calculated reflecting expressions relative to the mean of the chow group.

### High-resolution respirometry

After 48 hrs culture as described above but with additional electrical stimulation (C-pace EM, IonOptix, Westwood, MA, USA) with 1 Hz (10 Volt, 5 msec), Percoll gradient centrifugation was performed as described above for immunoblot and cell pellet was collected and washed three times with 4 ml mitochondrial respiration buffer MiR05 (60101-01, Oroboros, Innsbruck, Austria). The oxygen consumption rate (OCR) of the cardiomyocytes was measured using a polarographic oxygen sensor (O2K-fluo, Oroboros Instrument, Innsbruck, Austria) and a substrate-uncoupler-inhibitor titration protocol (SUIT). Briefly, 10,000 cells were added to the respiration chambers containing 2 ml MiR05 buffer, at 37 °C. The cells were first permeabilized by addition of 25 µg/ml saponin (Sigma Aldrich), followed by sequential addition of substrates (Sigma Aldrich) and inhibitors (Sigma Aldrich) of the mitochondrial electron transport chain: 0.5 mM carnitine, 40 µM palmitoyl-CoA, 0.1 mM malate, 1.5 mM Mg^2+^, 2.5 mM ADP, 5 mM pyruvate, 6 mM glutamate, 2 mM malate, 10 µM cytochrome c, 10 mM succinate, 2.5 µM oligomycin, 2 µM FCCP, 0.5 µM rotenone and 2.5 µM antimycin A.

### Immunocytochemistry

For immunofluorescence staining of isolated, cultured, purified cardiomyocytes, cells were fixed with 4% paraformaldehyde solution, permeabilized with 0,3% Triton X-100, and blocked with 10% normal goat serum. Antibodies to TOM70 (14528-1-AP, 1:200, Proteintech; Alexa Fluor 488-conjugated secondary antibody: A-11034, 1:500, Invitrogen), and troponin T (1:100, 565744, BD Pharmingen) were used, as well as DAPI (1:1000, ab228549, Abcam) for nuclei staining. Images were acquired with the BZ-X 810 microscope (Keyence) for densitometric quantification of TOM70.

### Statistical analysis

All data are shown as mean ± standard deviation (SD) unless otherwise indicated. The sample size is listed as N. Statistical differences were determined using GraphPad Prism Version 9.0 for Windows. Data were tested for normality using the Kolmogorov-Smirnoff test. For multiple independent groups, one-way analysis of variance (ANOVA) followed by Bonferronìs post-hoc test for parametric, and Kruskal-Wallis test followed by Dunn’s multi-comparison test for non-parametric data were used, unless otherwise indicated. For a within-group comparison over time, a marginal linear mixed effect model with Bonferronìs post-hoc test was used. Fixed variables were timepoint, treatment and interaction, while the individual animal was included as a random variable. Unless otherwise stated, the significances are shown compared to the respective control data. In detail, \**p* < 0.05; \*\**p* < 0.01; \*\*\**p* < 0.001.

## Results

### Nitro-oleic acid ameliorates diastolic dysfunction in mice with HFpEF

A mouse model of HFpEF induced by a combination of HFD and inhibition of eNOS with L-NAME for 5 or 15 weeks (wk) has been described and extensively characterized before by the group of Joseph Hill^25,32–34^. We chose the 15 wk HFD+L-NAME model and treated mice with NO_2_-OA (HFD+L-NAME NO_2_-OA) or vehicle (HFD+L-NAME vehicle) during wk 12 to 15 via osmotic pumps (Fig. 1a). Body weight and plasma glucose levels were markedly increased in HFD+L-NAME compared to mice fed with standard chow (control mice) (Fig. S1). As described for obese leptin-deficient mice^24^, treatment with NO_2_-OA significantly normalized glucose tolerance compared to vehicle. In addition, body weight was reduced (Fig. S1). Importantly, treatment with oleic acid (HFD+L-NAME OA) had no significant effect (Fig. S1), affirming the key impact of the nitroalkene substituent, which confers electrophilicity and promotes post-translational modification of predominantly cysteine residues. In line with the metabolic phenotype, analysis of isolated pancreatic islets revealed a rise in insulin content, indicative of compensatory islet enlargement in response to HFD+L-NAME. Furthermore, β-cell size was significantly increased. Similar trends were observed for islet- and α-cell size. Administration of NO_2_-OA normalized these parameters (Fig. S2).

**Fig 1:**
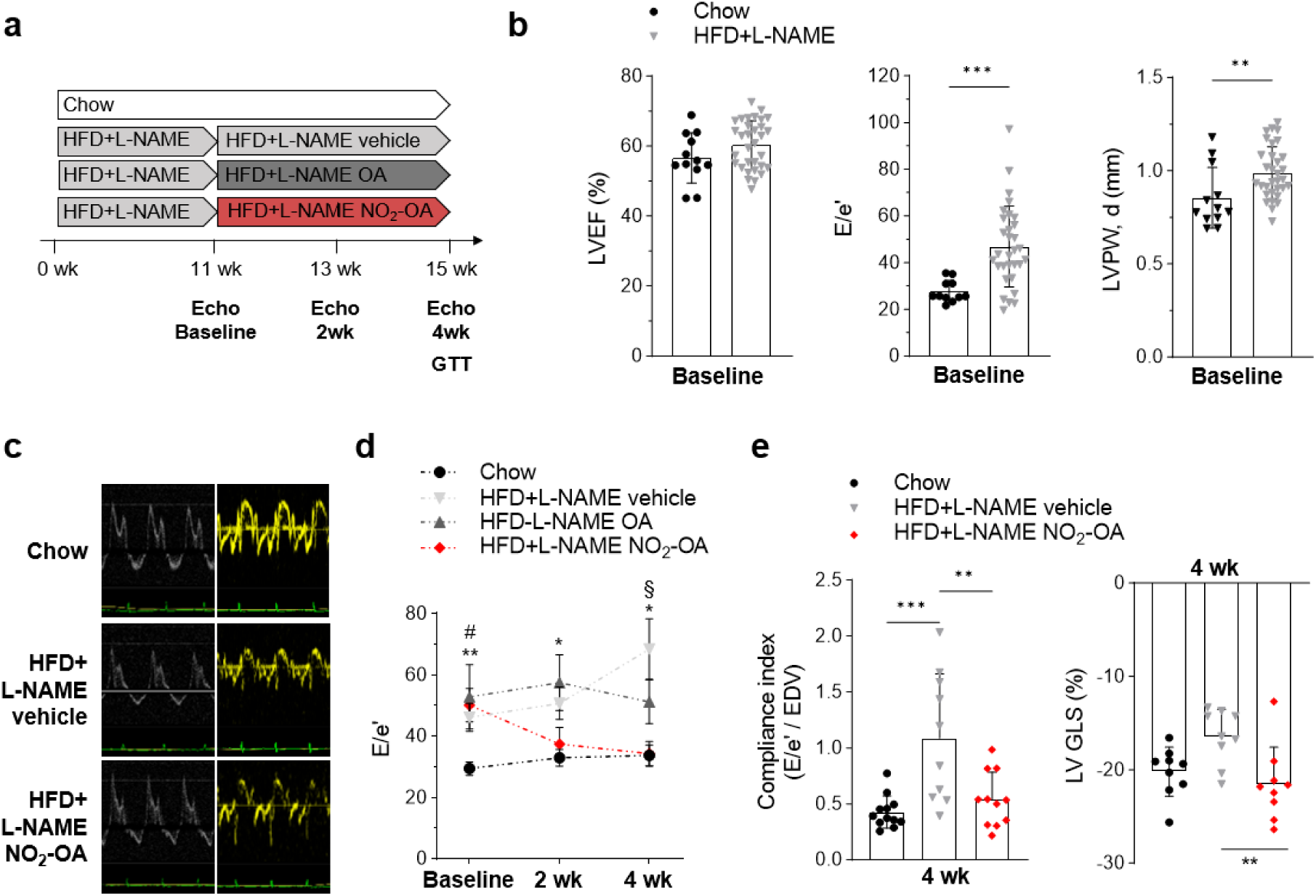
Nitro-oleic acid ameliorates diastolic dysfunction in mice with HFpEF. **a** Mice received high-fat diet and the endothelial nitric oxide synthase inhibitor L-NAME (HFD+L-NAME) or chow diet (Chow) for 15 weeks (wk) and were treated with vehicle-, oleic acid (OA)- or nitro oleic acid (NO_2_-OA) for the last 4 wk. Echocardiography (Echo) was performed after 11 wk of chow or HFD+L-NAME (Baseline), after 13 wk and 15 wk of chow or HFD+L-NAME including 2 wk or 4 wk of vehicle-, OA- or NO_2_-OA-treatment (2 wk, 4 wk). Glucose tolerance test was conducted at the final time point (GTT). **b** Left ventricular ejection fraction (LVEF), ratio of early diastolic mitral inflow velocity to early diastolic mitral annulus velocity (E/é) and left ventricular posterior wall thickness (LVPW, d) after 11 wk of chow or HFD+L-NAME (Baseline) (N=12/32). **c** Representative ultrasound images of LV pulse wave doppler (left) or tissue doppler at mitral annulus (right) to assess diastolic function. **d** Trend of E/é ratio during treatment of HFD+L-NAME mice with vehicle, OA or NO_2_-OA compared to chow mice (N=12/10/11/11). **e** Compliance index shown as E/é in ration to the individual end diastolic volume (EDV) (N=12/10/11) (left) and LV global longitudinal strain (GLS) (N=9/9/9) (right) after 15 wk of chow or HFD+L-NAME including 4 wk of vehicle- or NO_2_-OA treatment. Data are mean ± stardard deviation for **b** and **e**. **d** is shown as mean ± standard error of the mean. Statistical significance was calculated by unpaired, two-sided Student’s t-test for **b** and mixed effect analysis followed by Bonferronís multiple comparison test in **d**. * indicates statistical significance between chow and untreated HFD+L-NAME. # indicates statistical significance between chow and NO_2_-OA treated HFD+L-NAME. § indicates statistical significance between NO_2_-OA treated and untreated HFD+L-NAME. Statistical significance in **e** was calculated by One-way ANOVA followed by Bonferronís post-hoc test. *p*: **<0.01, ***<0.001.

To assess LV function, control and HFD+L-NAME mice before (baseline) and after 2 and 4 wk of treatment with NO_2_-OA, OA or vehicle were analysed by echocardiography (Fig. 1a). As previously^25^, HFD+L-NAME mice developed a HFpEF phenotype reflected by preserved LV ejection fraction (LVEF), diastolic dysfunction and LV hypertrophy at baseline (11 wk) (Fig. 1b) and interstitial fibrosis (Fig. S3a). Remarkably, treatment with NO_2_-OA significantly reduced the ratio of early diastolic mitral inflow velocity to early diastolic mitral annulus velocity (E/e’ ratio) compared to vehicle-treated HFD+L-NAME mice (Fig. 1c, d). The normalization of the early diastolic mitral annulus velocity (e’) after treatment with NO_2_-OA was primarily responsible for the improved diastolic function (Fig. S4), whereas the transmitral profile (E/A ratio) was not altered among the groups (Fig. S4). The E/e’ ratio related to end-diastolic volume reflecting LV compliance was also normalized by NO_2_-OA treatment (Fig. 1e). Importantly, the therapeutic effect of NO_2_-OA became also evident by a restoration of LV global longitudinal strain (Fig. 1e). In contrast, LV hypertrophy and fibrosis were not significantly influenced by NO_2_-OA (Fig. S3a, b). Taken together, treatment with NO_2_-OA eliminated diastolic dysfunction.

### Nitro-oleic acid restores protein profiles in LV of mice with HFpEF

To define the effects of NO_2_-OA treatment on the LV proteome, LV tissue of control, HFD+L-NAME vehicle and HFD+L-NAME NO_2_-OA mice were analysed by LC-MS (Fig. 2a). In total, 1796 proteins were identified. Unexpectedly, the principal component analysis indicated relatively slight differences in protein abundances between HFD+L-NAME and control mice, while HFD+L-NAME NO_2_-OA mice were clearly separated (Fig. 2b). In LV of HFD+L-NAME mice, only 1.3% of total proteins had significantly higher abundances and 1.7% of total proteins had lower levels compared to control (Fig. 2c). According to the lipid oversupply in vehicle-treated HFD+L-NAME mice, KEGG pathway analysis identified increased protein levels in cholesterol and lipid metabolism as well as redox signaling and lipid homeostasis in peroxisome in LV of HFD+L-NAME compared to control mice (Fig. 2d). Comparison of proteome data of NO_2_-OA versus vehicle-treated HFD+L-NAME mice showed, that 2.8% of total proteins had significantly lower levels, but 26.8% of total proteins had significantly increased abundance compared to vehicle (Fig. 2e), and 25.7% were significantly enhanced compared to both vehicle-treated as well as to chow mice. Further analyses of these proteins are shown below (Fig. 3-5). In addition, the LC-MS analysis showed that 70% of proteins (21 proteins) with decreased abundance in HFD+L-NAME compared to control hearts had a significantly higher abundance after NO_2_-OA compared to vehicle treatment (Fig. 2f). Thus, treatment with NO_2_-OA significantly normalized protein abundance after 15 wk of HFD+L-NAME administration.

**Fig. 2:**
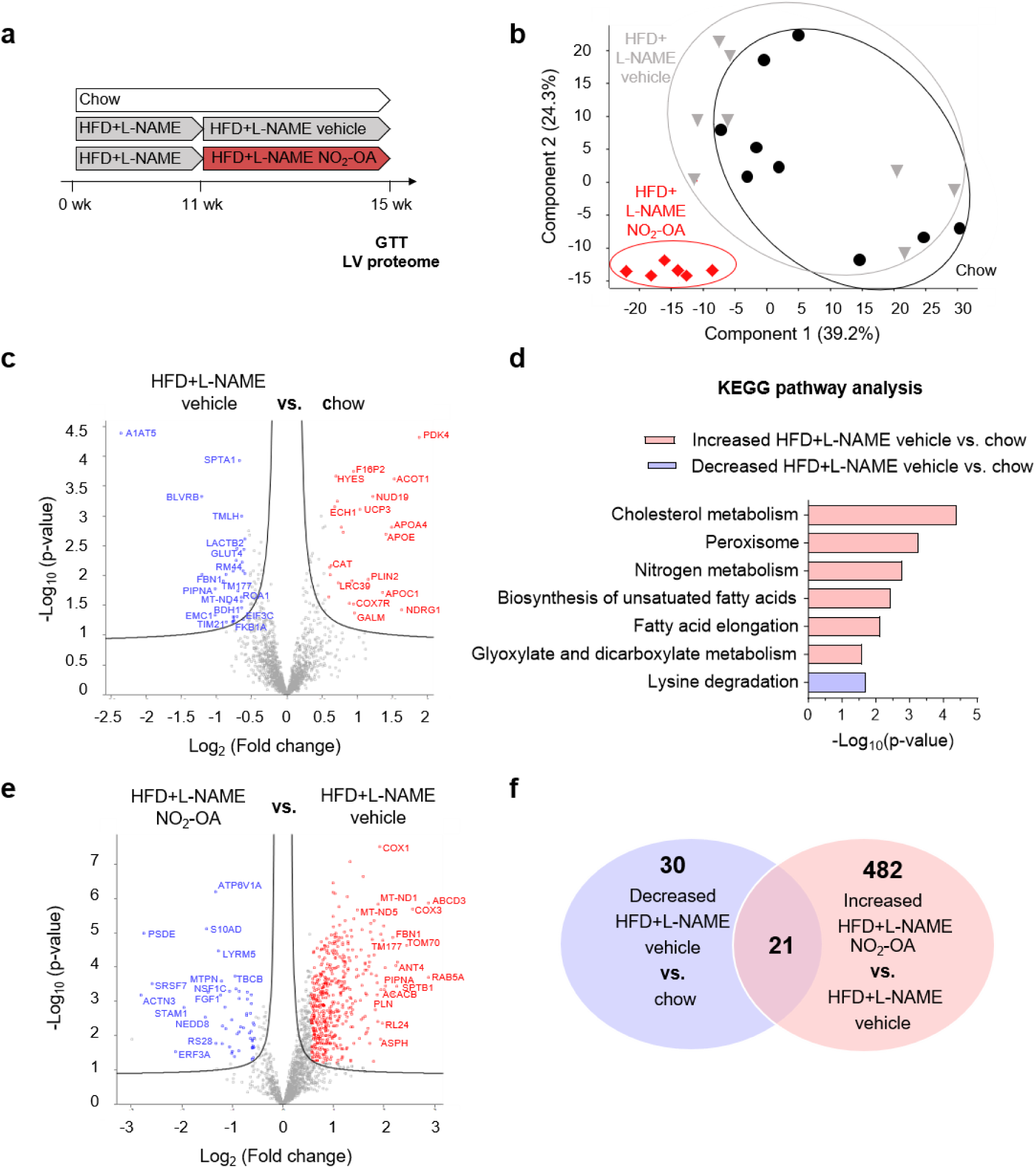
Myocardial proteome analysis in HFpEF mice reveals marked alterations upon nitro-oleic acid treatment. **a** Mice received high-fat diet and the endothelial nitric oxide synthase inhibitor L-NAME (HFD+L-NAME) or chow diet (Chow) for 15 weeks (wk) and were treated with nitro-oleic acid (NO_2_-OA) or vehicle for the last 4 wk. At the final time point proteome analysis of left ventricular tissue (LV proteome) was conducted. Glucose tolerance test (GTT) served as proof of effectiveness. **b** Principle component analysis of the proteome data set of LV mouse tissue (chow: N=9; HFD+L-NAME vehicle: N=8; HFD+L-NAME NO_2_-OA: N=6). **c, e** Volcano-plots of proteome data comparing HFD+L-NAME vehicle with chow mice (**c**) and NO_2_-OA-treated with vehicle-treated HFD+L-NAME mice (**d**). Significantly changed proteins are marked (significance ≥ - log_10_(0.05); red: fold change > log_2_(1.5); blue: fold change < log_2_(1.5)). **d** KEGG pathway enrichment analysis of significantly altered proteins of HFD+L-NAME vehicle compared to chow mice (protein count ≥ 2; red: analysis with increased proteins, blue: analysis with decreased proteins, significance of included proteins ≥ - log_10_(0.05)). **f** Venn diagram of proteins significantly decreased in the HFD+L-NAME vehicle compared to the chow group and proteins significantly increased after treatment with NO_2_-OA compared to untreated HFD+L-NAME vehicle mice. Statistical significance was calculated with Perseus (MPI Biochemistry) for **c** and **e** and with two-tailed Fisher’s exact test for **d**. *p*-values are shown as −log10(*p*-value). Abbreviations of proteins: A1AT5: alpha-1 antitrypsin; SPTA1: spectrin alpha; BLVRB: biliverdin reductase B; TMLH: trimethyllysine hydroxylase; LACTB2: lactamase beta 2; GLUT4: solute carrier family 2, member 4 (glucose transporter); RM44: mitochondrial ribosomal protein L44; FBN1: fibrilin 1; TM177: transmembrane protein 177; ROA1: heterogenous nuclear ribonucleoprotein A1; PIPNA: phosphatidylinositol transfer protein alpha; MT-ND4: NADH:ubiquinone oxidoreductase core subunit 4; BDH1: 3-hydroxybutyrate dehydrogenase 1; EIF3C: eukaryotic translation inhibition factor 3 subunit C; FKB1A: prolyl isomerase 1A; EMC1; endoplasmic reticulum membrane protein complex subunit 1; TIM21: translocase of inner mitochondrial membrane 21; PDK4: pyruvate dehydrogenase kinase 4; F16P2: fructose-bisphosphatase 2; ACOT1: acyl-CoA thioesterase 1; HYES: epoxide hydrolase 2; NUD19: nudix hydrolase 19; UCP3: uncoupling protein 3; ECH1: enoyl-CoA hydratase 1; APOA4: apolipoprotein A4; APOE: apolipoprotein E; CAT: catalase; PLN2: perilipin 2; LRC39: leucine rich repeat containing 39; APOC1: apolipoprotein C1; COX7R: cytochrome C oxidase subunit 7A2 like; NDRG1: N-myc downstream regulated 1; GALM: galactose mutarotase; ATP6V1A: ATPase H+ transporting V1 subunit A; S10AD: S100 calcium binding protein A13; PSDE: proteasome 26S subunit, non-ATPase 14; LYRM5: electron transfer flavoprotein regulatory factor 1; ACTN3: actinin alpha 3; SRSF7: serine and arginine rich splicing factor 7; MTPN: myotrophin; NSF1C: NSFL1 cofactor; FGF1: fibroblast growth factor 1; TBCB: tubulin folding cofactor B; STAM1: signal transducing adaptor molecule; NEDD8: ubiquitin like modifier; RS28: ribosomal protein S28; ERF3A: G1 to S phase transition 1; COX1: cytochrome c oxidase 1; MT-ND1; NADH:ubiquinone oxidoreductase core subunit 1; MT-ND5: NADH:ubiquinone oxidoreductase core subunit 5; ABCD3: ATP binding cassette subfamily D member 3; COX3: cytochrome c oxidase 3; TOM70: translocase of outer mitochondrial membrane 70; ANT4: adenine nucleotide translocator 4; SPTB1: spectrin beta; ACACB: acetyl-CoA carboxylase beta; PLN: phospholamban; RL24: ribosomal protein L24; ASPH: aspartate beta-hydrolase.

**Fig. 3:**
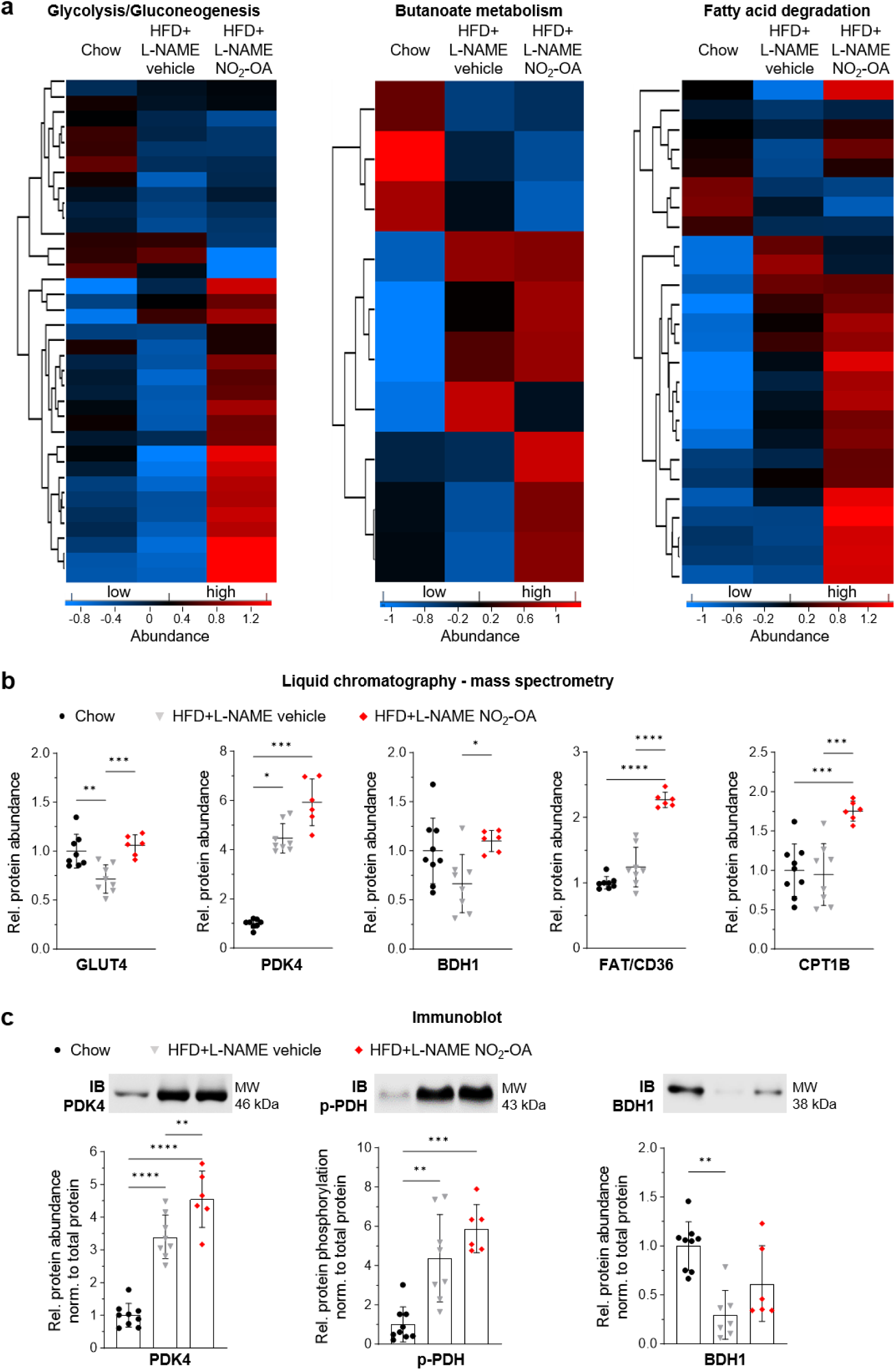
Nitro-oleic acid induces a beneficial metabolic protein profile in LV of mice with HFpEF. **a** Heatmaps presenting the protein abundance of all proteins associated with the indicated KEGG pathways for left ventricular (LV) tissue of mice after chow diet, 15 weeks of HFD and L-NAME with 4 weeks of vehicle (HFD+L-NAME vehicle) or NO_2_-OA treatment (HFD+L-NAME NO_2_-OA) obtained by liquid chromatography – mass spectrometry (LC-MS). **b** Relative protein abundance derived from LC-MS of GLUT4, PDK4, BDH1, FAT/CD36, CPT1B in LV mouse tissue of the different experimental groups. **c** Relative protein level normalized to total protein abundance and representative immunoblot (IB) for PDK4, p-PDH and BDH1 in LV mouse tissue of the different experimental groups. Protein levels were normalized to the total protein amount assessed on IB by fluorescence labelling (Fig. S7 and S8). Data in **b** and **c** are shown relative to the mean of the chow group. Statistical significance was calculated with One-way ANOVA followed by Bonferronís post-hoc test for **b** and **c**, except for PDK4 in **b**. Kruskal-Wallis test followed by Dunn’s multi-comparison test was used for PDK4 in **b**. N represent individual animals (N=9/8/6); outliers were identified by ROUT method and excluded from analysis. *p*: *<0.05, **<0.01, ***<0.001, ****<0.0001. Abbreviations of proteins: GLUT4: solute carrier family 2, member 4 (glucose transporter); PDK4: pyruvate dehydrogenase kinase 4; FAT/CD36: long chain fatty acid transporter; BDH1: 3-hydroxybutyrate dehydrogenase 1; CPT1B: carnitine O-palmitoyl-transferase 1, muscle isoform; p-PDH: phosphorylated pyruvate dehydrogenase at serine residue 293.

**Fig 4:**
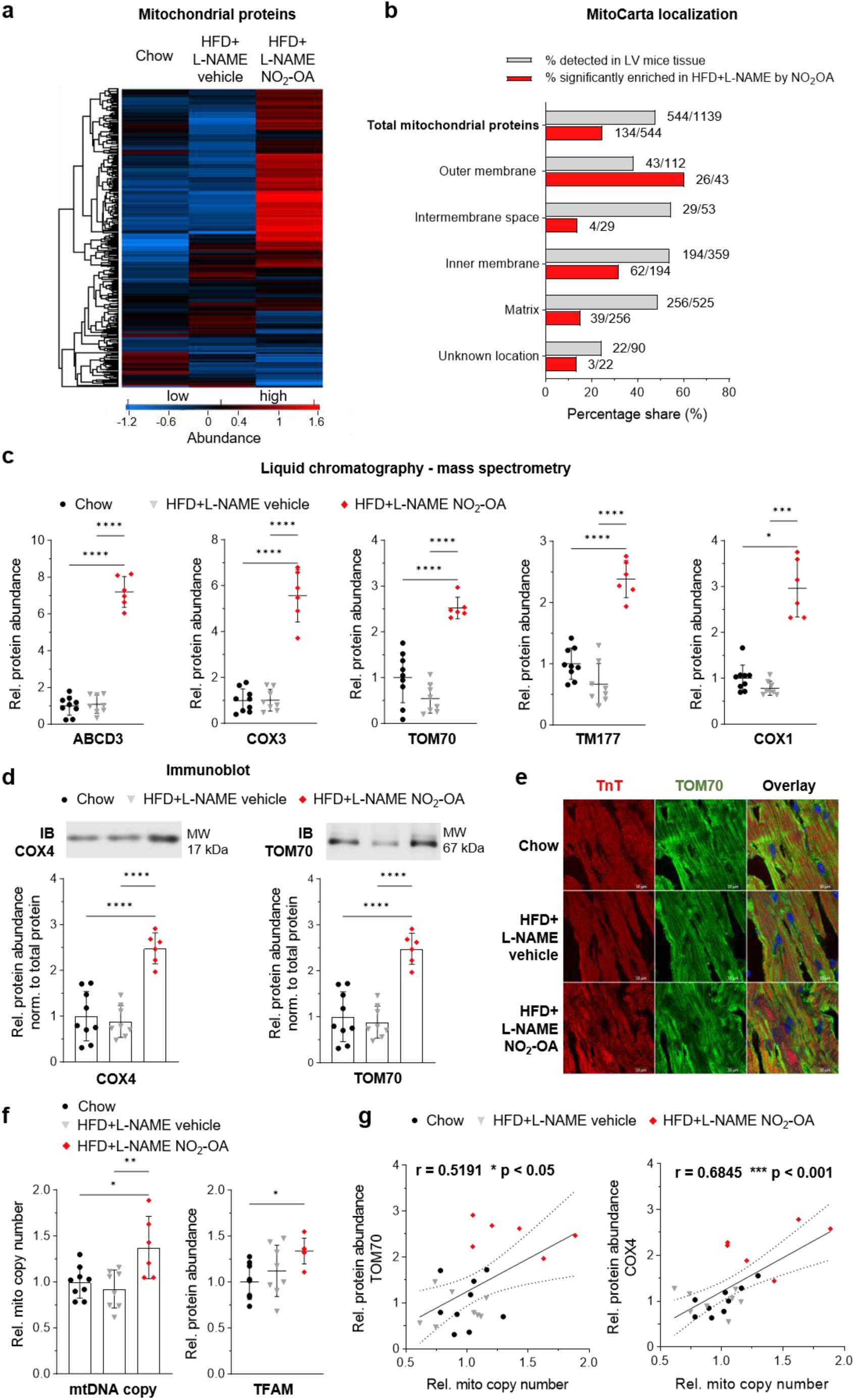
Mitochondrial biogenesis is stimulated by nitro-oleic acid in LV of mice with HFpEF. **a b** Identification of mitochondrial proteins (**a**) and localization (**b**) by annotation to the mitoCarta data base^37^ **a** Heatmaps presenting the protein abundance of all mitochondrial proteins detected by liquid chromatography – mass spectrometry (LC-MS) in left ventricular (LV) tissue of mice after chow diet, 15 weeks of HFD and L-NAME with 4 weeks of vehicle (HFD+L-NAME vehicle) or NO_2_-OA treatment (HFD+L-NAME NO_2_-OA). **b** Shown is the percentage of all mitochondrial proteins detected in LV mouse tissue by LC-MS (grey) and the percentage of mitochondrial proteins with significantly enhanced abundance after administration of NO_2_-OA in HFD+L-NAME treated mouse compared to HFD+L-NAME vehicle treatment (red). **c** Protein levels of the five most abundant mitochondrial proteins (ABCD3, COX3, TOM70, TM177 and COX1) in LV mouse tissue of the different experimental groups (-log_10_(p-value) > 4.5; log_2_(fold change) > 1.9). **d** Representative immunoblots (IB) and quantification of the mitochondrial proteins COX4 and TOM70 normalized to the total protein abundance of LV mouse tissue of the different experimental groups. Protein levels were normalized to the total protein amount assessed on IB by fluorescence labelling (Fig. S7 and S8). **e** Immunohistochemistry of TOM70 (green) and TnI (red) in LV mouse tissue of the three experimental groups. **f** Relative mitochondrial (mito) DNA (mtDNA) copy number und protein abundance of the mitochondrial transcription factor (TFAM) detected by LC-MS. **g** Correlation of mitochondrial protein level with relative mitochondrial copy number of LV mouse tissue in the different experimental groups. Data in **c, d** and **f** are shown relative to the mean of the chow group. Statistical significance was calculated with One-way ANOVA followed by Bonferronís post-hoc test for **c, d** and **f**. Pearson correlation was used for **g**. N represent individual animals (N=9/8/6). *p*: *<0.05, **<0.01, ***<0.001, ****<0.0001. Abbreviations of proteins: ABCD3: ATP binding cassette subfamily D member 3; COX3: cytochrome c oxidase 3; TOM70: translocase of outer mitochondrial membrane 70; TM177: transmembrane protein 177; COX1: cytochrome c oxidase 1; COX4: cytochrome c oxidase 4; TnI: troponin I; TFAM: mitochondrial transcription factor A.

**Fig 5:**
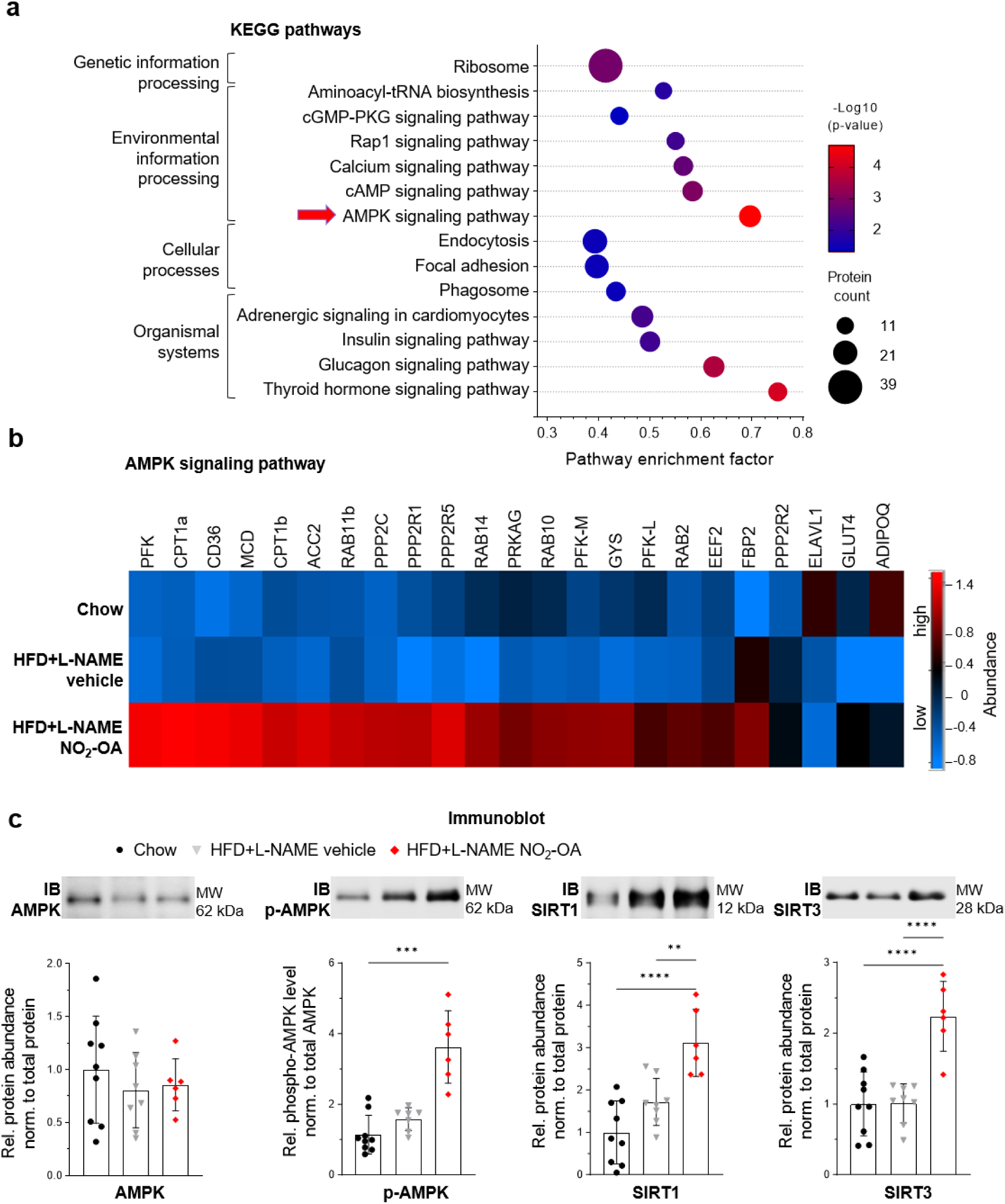
Activation of the AMPK signaling pathway by nitro-oleic acid in HFpEF mice hearts. **a** KEGG pathway enrichment analysis of proteins derived from liquid chromatography followed by mass spectrometry (LC-MS) with significantly enhanced abundance after administration of NO_2_-OA in HFD+L-NAME treated mouse hearts compared to vehicle treated mouse hearts (HFD+L-NAME vehicle) (inclusion criteria: protein count ≥ 10; pathway enrichment factor ≥ 30%; significance ≥ −log_10_(0.05)). The most significant KEGG pathway is marked with a red arrow. **b** Heatmap of all proteins associated with the AMPK signaling pathway and identified by LC-MS analysis in LV mouse tissue in the experimental groups. **c** Representative immunoblots and quantification of AMPK and p-AMPK, SIRT1 and SIRT3 of the different experimental groups. Protein levels were normalized to the total protein amount assessed on IB by fluorescence labelling (Fig. S7 and S9). Data in **c** are shown relative to the mean of the chow group. Statistical significance was calculated by two-tailed Fisher’s exact test for a and One-way ANOVA followed by Bonferronís post-hoc test for c. Kruskal-Wallis test with Dunńs multiple comparisons test was used for p-AMPK. N represent individual animals (N=9/8/6). *p*: **<0.01, ***<0.001, ****<0.0001. Abbreviations of proteins: PFK: phosphofructokinase; CPT1a, carnitine palmitoyltransferase 1A; CD36/FAT: long chain fatty acid transporter; MCD: malonyl-CoA decarboxylase; CPT1b: carnitine O-palmitoyltransferase 1B; ACC2: acetyl-CoA carboxylase beta; RAB11b: ras related protein 11b; PPP2C: serine/threonine-protein phosphatase 2A, catalytic subunit; PPP2R1: serine/threonine-protein phosphatase 2A, regulatory subunit A; PPP2R5: serine/threonine-protein phosphatase 2A, regulatory subunit B; PRKAG: 5’-AMP-activated protein kinase, regulatory gamma subunit; RAB10: ras related protein 10; PFK-M: phosphofructokinase, muscle isoform; GYS: glycogen synthase; PFK-L: phosphofructokinase, liver isoform; RAB2: ras related protein 2; EEF2: eukaryotic translation elongation factor 2; FBP2: fructose bisphosphatase 2; PPP2R2: serine/threonine-protein phosphatase 2A, regulatory subunit B; ELAVL1: ELAV-like protein 1; GLUT4: solute carrier family 2, member 4 (glucose transporter); ADIPOQ: adiponectin; AMPK: 5’-AMP-activated protein kinase; SIRT1/3, NAD-dependent protein deacetylase sirtuin-1/3; p-AMPK: phosphorylated 5’-AMP-activated protein kinase at threonine residue 172.

### Nitro-oleic acid alters the metabolic protein profile in LV of mice with HFpEF

As expected for the obesity-related HFpEF mouse model, the cardiometabolic protein profile was significantly impacted, as reflected by the heat maps showing changes in proteins associated with glycolysis/gluconeogenesis, butanoate metabolism and FA degradation (Fig. 3a). Cardiac protein levels of key enzymes of these pathways were also profoundly enhanced by the treatment of HFD+L-NAME mice with NO_2_-OA (Fig. 3b). In detail, protein abundance of the glucose transporter (GLUT4) was significantly decreased, and the pyruvate dehydrogenase kinase (PDK4) was significantly increased in HFD+L-NAME mice compared to control mice. Upon NO_2_-OA treatment, the GLUT4 level was normalized and the PDK4 level was further increased in HFD+L-NAME (Fig. 3b); these findings were validated using immunoblots (Fig. 3c). Accordingly, enhanced phosphorylation of pyruvate dehydrogenase (PDH) was detected in HFD+L-NAME mice, which was more profound in HFD+L-NAME NO_2_-OA (Fig. 3c), indicating an attenuation of GO rates. In line with previously published data of an HFpEF mouse model^35^, for butanoate metabolism the amount of D-beta-hydroxybutyrate dehydrogenase (BDH1) was lowered in HFD+L-NAME mice (indicating reduced ketone body utilization) and was normalized by NO_2_-OA (Fig. 3b, LC-MS; Fig. 3c, immunoblotting). Both effects, the normalization of ketolysis and the further decrease in GO by NO_2_-OA might contribute to improved substrate utilization, energy supply, and cardioprotection^36^. This is supported by the significant effects of NO_2_-OA on mediators of FAO. Whereas subtle changes were detected in LV of HFD+L-NAME mice compared with control mice, NO_2_-OA induced significant elevations in proteins responsible for lipid handling, particularly concerning the levels of fatty acid translocase (FAT, CD36) and carnitine palmitoyltransferase 1B (CPT1B) (Fig. 3b). This indicates augmented FA uptake and metabolism, which might result not only in enhanced energy generation, but also in the mitigation of lipotoxicity.

### Nitro-oleic acid increases mitochondrial proteins in LV of mice with HFpEF

With LC-MS analysis, 544 out of 1139 mitochondrial proteins annotated in the mitoCarta database^37^ were detected. Remarkably, in LV of NO_2_-OA-treated mice, 24.6% of these mitochondrial proteins were significantly more abundant than in vehicle-treated mice, which is impressively demonstrated by the heatmap (Fig. 4a). MitoCarta localization^37^ illustrates the functional affiliation of these mitochondrial proteins and the percentage of those significantly increased by NO_2_-OA treatment (Fig. 4b). This analysis revealed that mitochondrial outer membrane proteins are particularly enriched by NO_2_-OA (Fig. 4b). Fig. 4c illustrates the 5 most abundant mitochondrial proteins identified in NO_2_-OA treated HFD+L-NAME mouse hearts by LC-MS compared to vehicle-treated HFD+L-NAME mice, emphasizing the significant upregulation of these mitochondrial proteins by NO_2_-OA. This was confirmed in immunoblots of the mitochondrial proteins cytochrome c oxidase subunit 4 (COX4) and the mitochondrial transport protein TOM70 (Fig. 4d), as reflected by immunohistochemistry of TOM70 protein in LV tissue sections (Fig. 4e). In contrast, the *TOMM70* mRNA level in LV was unchanged among the groups (Fig. S5). Interestingly, from the 13 mitochondrially encoded proteins, 8 were initially detected by LC-MS and shown to be significantly enhanced after treatment with NO_2_-OA in HFD+L-NAME compared to control and vehicle. Of note, mitochondrial transcription factor A (TFAM) was significantly enhanced by NO_2_-OA in HFD+L-NAME mice compared to chow and HFD+L-NAME mice (Fig. 4f), pointing towards increased mitochondrial biogenesis. In line, mitochondrial DNA copy number was significantly increased in these animals (Fig. 4f) and the relative number of mitochondrial DNA copies strongly correlated with the protein expression of TOM70 and COX4 (Fig. 4g). Taken together, the proteome analysis, immunoblot, and immunohistochemistry of LV tissue showed an increase in mitochondrial proteins in NO_2_-OA-treated mice.

### Nitro-oleic acid upregulates sirtuin expression via induction of AMPK signaling

KEGG pathway analysis of protein expression changes by NO_2_-OA identified AMPK signaling, an important mediator of mitochondrial biogenesis, as among the most significantly enriched pathways (Fig. 5a). The LC-MS heatmap of proteins associated with AMPK signaling illustrates a significantly greater protein abundance after treatment with NO_2_-OA in HFD+L-NAME mouse hearts compared to chow and HFD+L-NAME mice (Fig. 5b). Immunoblot analysis revealed significantly greater AMPK phosphorylation at threonine-172 in NO_2_-OA-treated compared to vehicle-treated HFD+L-NAME animals as well as compared to controls, whereas AMPK protein levels were not different between groups (Fig. 5c). The activation of AMPK resulted in significantly increased mRNA expression of the NAD^+^-dependent deacetylase sirtuin 1 (*Sirt1*), as previously^38^ (Fig. S6a). Also, SIRT1 protein levels were significantly enhanced by NO_2_-OA in HFD+L-NAME mouse hearts (Fig. 5c). SIRT1 is located in the cell nucleus and mediates gene transcription by deacetylation of transcription factors, such as the master regulator of mitochondrial biogenesis, peroxisome proliferator-activated receptor gamma coactivator 1-alpha (PGC-1α)^39,40^. In line with this, an increase of the mitochondrial deacetylase sirtuin 3 (SIRT3) expression was detected after treatment with NO_2_-OA, as compared to chow and HFD+L-NAME mice (Fig. 5c). Sirtuin activity might also be further induced by elevated NAD^+^ levels after treatment with NO_2_-OA, since there were significantly increased mRNA levels of nicotinamide nucleotide adenyl transferase 1 (*Nmnat1*) (Fig. S6b), an enzyme central to the biosynthesis of NAD^+^.

### Nitro-oleic acid increases metabolically-stressed cardiomyocyte mitochondrial respiration

To analyse the effects of NO_2_-OA on mitochondrial respiration, isolated adult murine cardiomyocytes were cultivated for 48 hrs under normal glucose conditions (Ctrl) and metabolic stress conditions induced by high glucose levels, endothelin-1 (ET-1) and hydrocortisone (HC) as described before^29^, with either methanol as a vehicle or NO_2_-OA (Fig. 6a). Immunoblot and immunocytochemical analysis (Fig. 6b-d) revealed enhanced mitochondrial protein expression accompanied by modest activation of AMPK by NO_2_-OA in metabolically stressed hyperglycaemic cardiomyocytes. High-resolution respirometry of cardiomyocytes showed significantly increased oxidative phosphorylation (OXPHOS) after treatment with NO_2_-OA (Fig. 6e, f).

**Fig 6:**
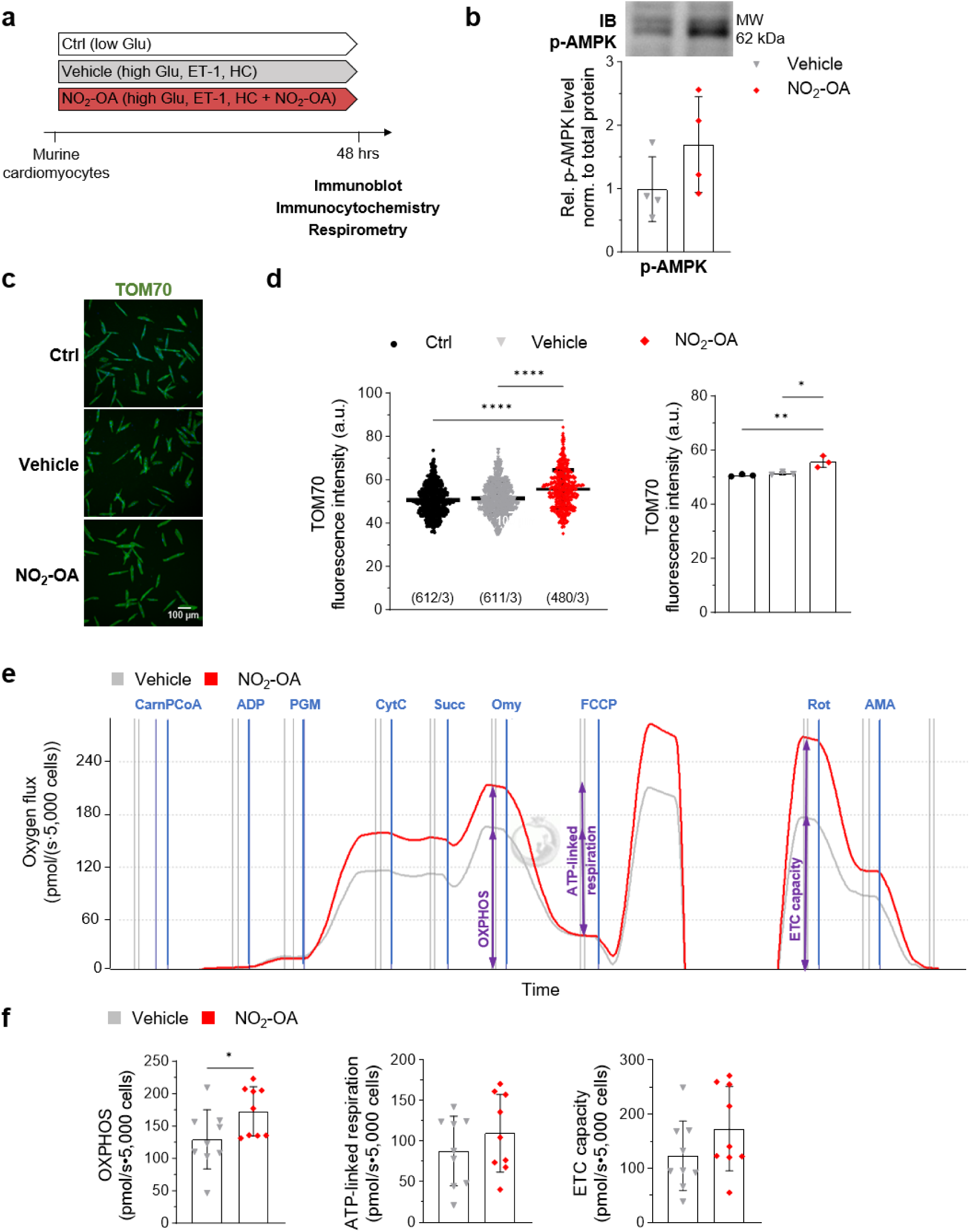
Nitro-oleic acid enhances mitochondrial function in metabolically-stressed cardiomyocytes. **a** Isolated adult murine cardiomyocytes were cultivated for 48 hrs under control conditions (Ctrl) with normal glucose levels (low Glu), under metabolic stress conditions induced by high glucose levels (high Glu), endothelin-1 (ET-1) and hydrocortisone (HC) with treatment of methanol (Vehicle) or nitro-oleic acid (NO_2_-OA). **b** Representative immunoblot and quantification of p-AMPK. Protein levels were normalized to the total protein amount assessed on IB by fluorescence labelling (Fig. S10) and shown relative to the individual Ctrl group (N=4). **c** Immunohistochemistry of TOM70 (green) in cardiomyocytes of the three experimental groups. **d** Quantification of TOM70 fluorescence intensity shown as arbitrary units (a.u.) for all analysed individual cardiomyocytes of three experiments (left) and as mean per experiment (right) (N=3). **e, f** High resolution respirometry of cardiomyocytes cultivated under metabolic stress conditions (high Glu, ET-1, HC) treated with methanol (Vehicle) or NO_2_-OA (NO_2_-OA) for 48 hrs. **e** Representative trace of oxygen flux for the two experimental groups. Addition of components are marked with blue lines (CarnPCoA: carnitine palmitoyl-CoA; ADP: adenosine diphosphate; PGM: pyruvate glutamate malate; CytC: cytochrome C; Succ: succinate; Omy: oligomycin; FCCP: mitochondrial uncoupler; Rot: rotenone; AMA: antimycin A). **e** Oxidative phosphorylation (OXPHOS), ATP-linked respiration and electron transport chain (ETC) capacity extracted from oxygen flux traces (N=9). Statistical significance was calculated by One-way ANOVA followed by Bonferronís post-hoc test for **d** and unpaired, two-sided Student’s t-test for **f**. *p*: *<0.05, **<0.01, ****<0.0001. Abbreviations of proteins: p-AMPK: phosphorylated 5’-AMP-activated protein kinase at threonine residue 172; TOM70: translocase of outer mitochondrial membrane 70.

## Discussion

Our study reveals that the small molecule electrophile NO_2_-OA significantly improved diastolic function in a ‘two-hit’ HFpEF mouse model. This effect was based on a marked alteration in mitochondrial and metabolic protein levels in the LV of NO_2_-OA-treated mice. The abundance of mitochondrial proteins including proteins responsible for lipid handling and FAO were significantly increased in LV tissue of NO_2_-OA-treated mice. These alterations were regulated by the activation of AMPK, which was detected in HFD+L-NAME NO_2_-OA mice and isolated cardiomyocytes treated with NO_2_-OA. AMPK is central to the regulation of cellular catabolic pathways, thereby inducing FAO and mitochondrial biogenesis^41^. LC-MS and immunoblot analyses of LV tissues indicated a significant increase in the abundance of AMPK-regulated proteins in NO_2_-OA-treated mice. Various signaling effects of NO_2_-OA were previously reported to be linked to changes in protein function through the reversible covalent nitro-alkylation of nucleophilic residues, primarily cysteine^42^. The deacetylase SIRT6 is activated by NO_2_-OA upon nitroalkene adduct formation^43^. Here, we found SIRT1 and SIRT3 protein levels increased in NO_2_-OA-treated mice, which are downstream targets of AMPK. Of note, SIRT1-induced activation of AMPK and positive feedback mechanisms between AMPK activation and SIRT1 induction^44^ as well as compensatory effects of SIRT1 and SIRT3 protein expression^45,46^ have been reported. It remains to be defined what mechanisms account for AMPK and SIRT pathway modulation by NO_2_-OA. PPARα agonism by NO_2_-OA might be of relevance, given the important role of PPARα in stimulating FAO in the heart, however, the agonistic effect of NO_2_-OA on PPARγ is stronger than that for PPARα^26^. Systemic PPARγ activation is likely to account for the anti-hyperglycaemic effect in the NO_2_-OA-treated mice, as previously^24^. However, the improvement in glucose tolerance is unlikely to be responsible for reducing diastolic dysfunction. Indeed, clinical data shows that the normalization of blood glucose in diabetic patients does not reduce cardiovascular death^47^, and thiazolidinedione PPARγ agonists increase the risk of heart failure despite effective blood glucose reduction in diabetic patients^48,49^. Instead, our data suggest that the NO_2_-OA-treated mice are characterized by a profound enhancement of mitochondrial biogenesis, with AMPK activation, increased TOM70 protein levels and increased OXPHOS in murine cardiomyocytes exposed to hyperglycaemic metabolic stress during electrical pacing. Small changes in some parameters might be related to the short-term treatment of 48 hrs due to the challenges of the long-term culture of isolated adult cardiomyocytes. Nonetheless, the results reveal a direct effect of NO_2_-OA on cardiomyocytes under conditions established for modeling diabetic cardiomyopathy^29^.

Given that in metabolic syndrome, FAO is increased due to lipid oversupply and resulting insulin resistance, it has been hypothesized that a reduction of FAO is beneficial in cardiometabolic HFpEF. However, this hypothesis has been disproven in experimental and clinical studies. In HFD-fed mice with inducible deletion of acetyl coenzyme A carboxylase 2, enhanced FAO protected from the development of HFpEF^50^. In this regard, the DoPING-HFpEF trial showed that inhibition of FAO with trimetazidine in HFpEF patients did not improve postcapillary pulmonary hypertension or exercise capacity^51^. This may in part be due to the fact that FA are highly energy efficient substrates. It has become evident that ATP deficiency is an important pathomechanistic characteristic of HFpEF. In fact, resting ATP levels are reduced to 20-40% in HFpEF^52–54^, and decreased phospho-creatine/ATP ratio correlates with diastolic dysfunction in obese HFpEF patients^55^. Given that the sarcoplasmic reticulum calcium ATPase (SERCA) is the most energy-demanding enzyme in the contractile apparatus, an energy deficit likely accounts for impaired relaxation^56^. Therefore, therapeutic strategies that increase energy yields hold promise. For example, SGLT2 inhibitors have been proposed to exert physiological benefit by inducing ketone metabolism^57^. Likewise, metformin, an AMPK activator, exerts cardioprotective effects in HFpEF^58^. Given that AMPK induces FAO and mitochondrial biogenesis, not only ATP production rates of cardiomyocytes are increased, but also the metabolism of high levels of FAs are enhanced and thus lipotoxicity is reduced. These reactions can promote the improvement of myocardial bioenergetics, thus restoring cardiomyocyte relaxation, as reflected by increased early diastolic mitral annulus velocity and global longitudinal strain. Together with NO_2_-OA’s already known advantageous effects in cardiac and metabolic disorders, this molecule possesses a unique repertoire of actions making it potentially relevant for clinical HFpEF therapy.

### Study limitations

This preclinical study has limitations. Most importantly, we have not determined ATP levels specifically in the LV of mice, nor have we assessed the rate of OXPHOS or FAO in this compartment. We do not know whether the increased abundance of LV mitochondrial proteins relate to increased metabolism, thus leading to the reliance on data from isolated cardiomyocytes treated *in vitro* with NO_2_-OA that indicated enhanced OXPHOS. Second, we have not determined lipid levels in plasma or myocardium. Therefore, defining changes in lipid handling and its effect on myocardial lipotoxicity is a future goal. Furthermore, we have not analysed calcium handling, which is important to understand the diastolic dysfunctional phenotype fully. Another important point is that we have not measured blood pressure in the mice. It has been previously reported that NO_2_-OA inhibits angiotensin II (AngII) type 1 receptor (AT1R) signaling thereby reducing blood pressure in AngII-treated mice^59^. Furthermore, NO_2_-OA inhibits soluble epoxide hydrolase, which leads to epoxyeicosatrienoic acid-mediated lowering of blood pressure^60^. Whether these mechanisms are of relevance in the present model remains elusive. Apart from that, the multi-target signaling actions of NO_2_-OA can exert protection from diastolic dysfunction in this model, i.e., enhanced activation of the ryanodine receptor^61^, modulation of mitochondrial function by nitroalkylation of uncoupling proteins (UCP) and adenine nucleotide translocase (ANT)^62,63^, improvement of vascular function via anti-inflammatory mechanisms and other potential actions.

### Conclusions and perspectives

Our study has identified enhanced mitochondrial biogenesis to be critical for restoring myocardial relaxation in a murine model of obese HFpEF. The small molecule nitroalkene NO_2_-OA induced these effects, which were associated with activation of AMPK signaling. Given the limited treatment options for HFpEF, these findings underline the promising biological properties of NO_2_-OA. An ongoing phase 2 trial testing NO_2_-OA in obese asthmatics (NCT03762395) should be helpful in developing further insight into the gene expression, metabolomic, inflammatory signaling and physiological responses that this small molecule nitroalkene can induce in obese subjects.

Despite the promising effects of SGLT2 inhibitors in HFpEF patients^5^ and the impressive effects of GLP-1 agonists on symptom burden in obese HFpEF^64^, more treatment options are warranted for this highly prevalent and morbid disorder, especially considering the limited tolerability to GLP-1 agonists^65^. With a positive safety and tolerability profile, NO_2_-OA emerges as a potentially important drug for treating HFpEF and other cardiopulmonary diseases^66^.

## Acknowledgements

We thank Désirée Gerdes and André Grafe for expert technical assistance.

## Author Contribution

All authors made substantial contributions to the conception or design of the study; the acquisition, analysis, or interpretation of data; or drafting or revising the paper. All authors approved the paper. All authors agree to be personally accountable for individual contributions and to ensure that questions related to the accuracy or integrity of any part of the work are appropriately investigated, resolved, and the resolution documented in the literature. M.M., U.S., A.K. conception and design of research; T.S., T.J.S., T.P., T.M., L.A.L., L.L., J.H., S.L., E.T., J.D., F.L.H. performed experiments; M.M., T.S., T.J.S., T.P., T.M., L.A.L., L.L., J.H., S.L., E.T., J.D., F.L.H., J.C.R., U.S., A.K. analysed data; M.M., T.S., C.W., B.S., M.D., E.T., J.D., S.H., K.L., J.R., U.S:, A.K. interpreted results of experiments; M.M., T.S., J.H., M.D., U.S., A.K. prepared figures; M.M., A.K. drafted the manuscript; M.D., S.H., K.L., F.S., B.F., V.R. edited and revised the manuscript.

## Supplemental Material

Expanded Materials and Methods

Supplemental Figures S1-S10

## Non-standard Abbreviations and Acronyms

4APIX: apical four chamber view
AMPK: 5’ adenosine monophosphate-activated protein kinase
AngII: angiotensin 2
ANT: adenosine nucleotide translocase
AT1R: angiotensin 2 type 1 receptor
ATP: adenosine triphosphate
AUC: area under the curve
BDH1: 3-hydroxybutyrate dehydrogenase 1
COX4: cytochrome c oxidase subunit 4
CPT1B: carnitine O-palmitoyl-transferase 1
DAG: diacylglycerol
eNOS: endothelial nitric oxide synthase
ET-1: endothelin-1
FA: fatty acid
FAO: fatty acid oxidation
FAT/CD36: long chain fatty acid transporter
gDNA: genomic DNA
GLUT4: glucose transporter, solute carrier family 2, member 4
GO: glucose oxidation
HC: hydrocortisone
HFD: high fat diet
HFpEF: heart failure with preserved ejection fraction
KEGG: Kyoto Encyclopedia of Genes and Genomes
LC-MS: liquid chromatography followed by mass spectrometry
L-NAME: *N*_ω_-nitro-L-arginine methyl ester hydrochloride
LV: left ventricle
LVEF: left ventricular ejection fraction
LVPW: left ventricular posterior wall thickness
mtDNA: mitochondrial DNA
NAD: nicotinamide adenine dinucleotide
NF-κB: nuclear factor ‘kappa-light-chain-enhancer’ of activated B-cells
*Nmnat1*: gene of nicotinamide nucleotide adenyl transferase 1
NO_2_-CLA: conjugated nitro-linoleic acid
NO_2_-LA: nitro-linoleic acid
NO_2_-OA: nitro-oleic acid
*Nos2*: gene of nitric oxide synthase, inducible
Nrf2: nuclear factor erythroid 2-related factor 2
OA: oleic acid
OCR: oxygen consumption rate
OXPHOS: oxidative phosphorylation
PDK4: pyruvate dehydrogenase kinase 4
PDH: pyruvate dehydrogenase
PGC-1α: peroxisome proliferator-activated receptor gamma coactivator 1-alpha PPAR peroxisome-proliferator activated receptor
PSAX: parasternal short-axis
PSLAX: parasternal long-axis
RPL32: ribosomal protein L32
SD: standard deviation
SEM: standard error of the mean
SERCA: sarcoplasmic reticular calcium ATPase
SGLT2: sodium glucose cotransporter 2
SIRT: sirtuin
*Sirt 1*: gene of sirtuin 1
SUIT: substrate-uncoupler-inhibitor titration protocol
TnT: troponin T
TOM70: translocase of outer membrane 70
*Tomm70*: gene of translocase of outer membrane 70
UCP: uncoupling protein
*Xbp1s*: gene of spliced form of X-box binding protein 1

