## Supplements for "Enhanced cardiac mitochondrial biogenesis by nitro-oleic acid remedies diastolic dysfunction in a mouse model of heart failure with preserved ejection fraction"

**Expanded Materials and Methods**

For cardiomyocyte isolation, the following buffers were used:

Perfusion buffer (pH 7,40, 20°C)

| | MW/ g/mol | $C_{final}$/ mM |
|---|---|---|
| NaCl | 58,4 | 113 |
| KCl | 74,6 | 4,7 |
| $Na_2HPO_4$ $2H_2O$ | 177,9 | 0,6 |
| $KH_2PO_4$ | 136,1 | 0,6 |
| $NaHCO_3$ | 84,0 | 12 |
| $KHCO_3$ | 100,1 | 10 |
| $MgSO_4$ $7H_2O$ | 246,5 | 1,2 |
| HEPES | 238,3 | 10 |
| Taurine | 125,1 | 30 |
| BDM | 101,1 | 10 |
| Glucose | 180,2 | 10 |

Digestion buffer

| | $C_{final}$ |
|---|---|
| Liberase TM | 100 µg/ml |
| $CaCl_2$ | 0,0125 mM |
| Perfusion buffer | |

Stop buffers

| | SB1 | SB2 | SB3 | SB4 |
|---|---|---|---|---|
| **$Ca^{2+}$-concentration/ mM** | **0,075** | **0,225** | **0,600** | **1,500** |
| Stop A | 95 % | 85 % | 60 % | 0 % |
| Stop B | 5 % | 15 % | 40 % | 100 % |

Stop A

| | $C_{final}$ |
|---|---|
| 10 % BSA-Solution | 1 % |
| Perfusion buffer | |

Stop B

| | $C_{final}$ |
|---|---|
| 100 % BCS | 10 % |
| M199 (Stock) | |

M199 (Stock)

| | $C_{final}$/ mM |
|---|---|
| Creatine-Monohydrate | 5 |
| L-Carnitine | 2 |
| Taurine | 5 |
| Pen/Strep | 100 U/ml |
| M199 (Hanks Salt) | |

#### Supplemental Figures

### Figure S1

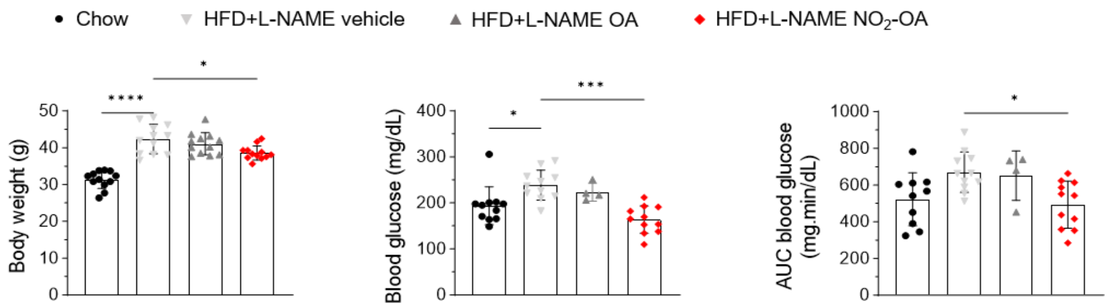


**Figure S1: Nitro-oleic acid reduces bodyweight and improves glucose tolerance in HFpEF mice.** Mice received high fat diet (HFD) and the endothelial nitric oxide synthase inhibitor L-NAME or chow diet (chow) for 15 weeks (wk) and were treated with oleic acid (OA) or nitro-oleic acid ($NO_2$-OA) or vehicle for 4 wk. Body weight (BW) (N=12/12/12/12), blood glucose levels during an intraperitoneal glucose tolerance test shown as blood glucose concentrations 90 min after glucose injection (N=11/11/4/11) and area under the curve (AUC) over the total time of 90 min (N=10/11/4/11). Statistical significance was calculated with One-way ANOVA followed by Bonferroni´s post-hoc test for BW and AUC. Kruskal-Wallis test followed by Dunn´s multiple comparisons test was used for raw blood glucose values. N represent individual animals. Variation in N is due to failed blood collection. Outliers were identified using ROUT method and excluded from analysis.

### Figure S2

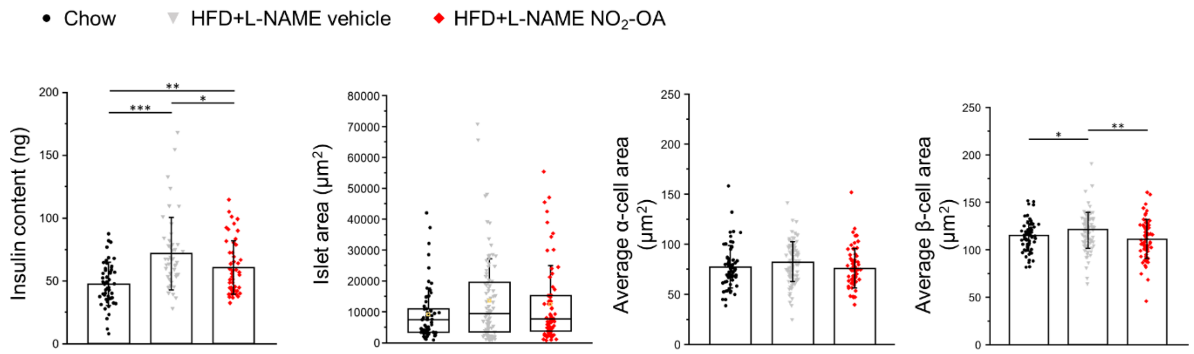


**Figure S2: Nitro-oleic acid partly counteracts the changes of the endocrine pancreas induced by HFD+L-NAME.**

Mice received high fat diet (HFD) and the endothelial nitric oxide synthase inhibitor L-NAME or chow diet (chow) for 15 weeks (wk) and were treated with nitro-oleic acid ($NO_2$-OA) or vehicle for 4 wk. Insulin content was determined per islet (10/10/10 animals per cohort) and analysis of whole islet and the average α- or β-cell size was performed in pancreatic tissue slices. Parameters are calculated as area per islet and cell, respectively (5/5/4 animals per cohort). One-way ANOVA followed by Neumann-Keuls post-hoc test was used for insulin content and the average cell area of α and β-cells. Kruskal-Wallis test followed by Dunn´s multiple comparisons was used for the islet area. N represent islets.

### Figure S3

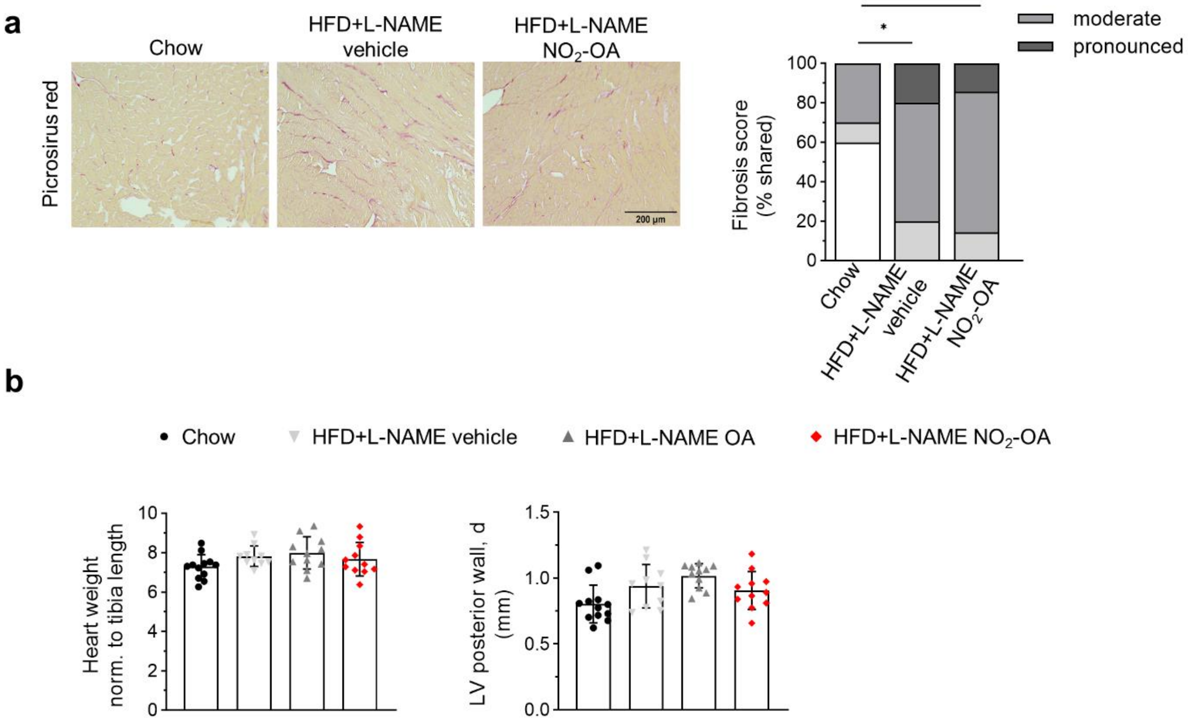


**Figure S3: Nitro-oleic acid has no effect on the structural remodeling in HFpEF mice.**

Mice received high fat diet (HFD) and the endothelial nitric oxide synthase inhibitor L-NAME or chow diet (chow) for 15 weeks (wk) and were treated with oleic acid (OA) or nitro-oleic acid ($NO_2$-OA) or vehicle for 4 wk. **a** Representative images (scale bar 200 µm) of left ventricular (LV) sections stained with picrosirius red and analysis of fibrosis grades (N=10/10/7). **b** Heart weight and LV posterior wall thickness as assessed by echocardiography is shown (N=12/10/11/11). Statistical significance was calculated by Fisher exact test for **a** and One-way ANOVA followed by Bonferroni´s post-hoc test for heart weight and Kruskal-Wallis test followed by Dunn´s multiple comparisons was used for LV posterior wall thickness **b**. * individual analysis comparing chow and HFD+L-NAME vehicle group. # individual analysis comparing HFD+L-NAME vehicle and $NO_2$-OA group. N represent individual animals.

### Figure S4

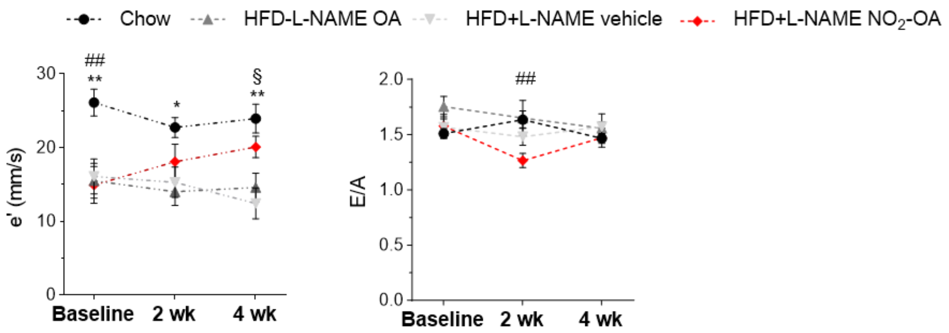


**Figure S4: Diastolic function in nitro-oleic acid-treated HFpEF mice.**

Echocardiography was performed in mice after 11 weeks (wk) of chow or high fat diet (HFD) and the endothelial nitric oxide synthase inhibitor L-NAME (Baseline), after 13 wk of chow or HFD and L-NAME including 2 wk of vehicle-, oleic acid (OA)- or nitro oleic acid ($NO_2$-OA)-treatment (2 wk) and after 15 wk of chow or HFD+L-NAME including 4 wk of vehicle-, OA- or $NO_2$-OA-treatment (4 wk) (N=12/10/11/11). Early diastolic mitral annulus velocity (e') was assessed by echocardiography using tissue doppler. The transmitral profile (E/A ratio) was assessed by pulse wave doppler. Statistical significance was calculated by mixed effect analysis followed by Bonferroni´s multiple comparison test. * indicates statistical significance between chow and untreated HFD+L-NAME. # indicates statistical significance between chow and $NO_2$-OA treated HFD+L-NAME. § indicates statistical significance between $NO_2$-OA treated and untreated HFD+L-NAME.

### Figure S5

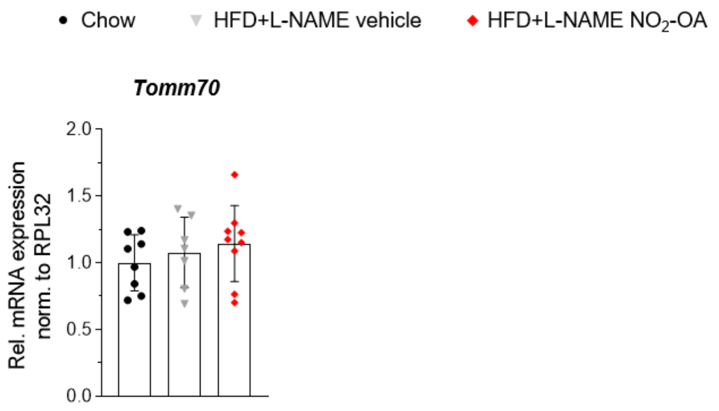


**Figure S5: *TOMM70* mRNA expression is not changed in LV tissue of HFpEF mice.**
Mice received high fat diet (HFD) and the endothelial nitric oxide synthase inhibitor L-NAME or chow diet (chow) for 15 weeks (wk) and were treated with nitro-oleic acid ($NO_2$-OA) or vehicle for 4 wk. mRNA expression of the translocase of outer mitochondrial membrane protein 70 (TOM70, gene: *Tomm70*) was performed in left ventricular mouse tissue (N=9/8/6).

### Figure S6

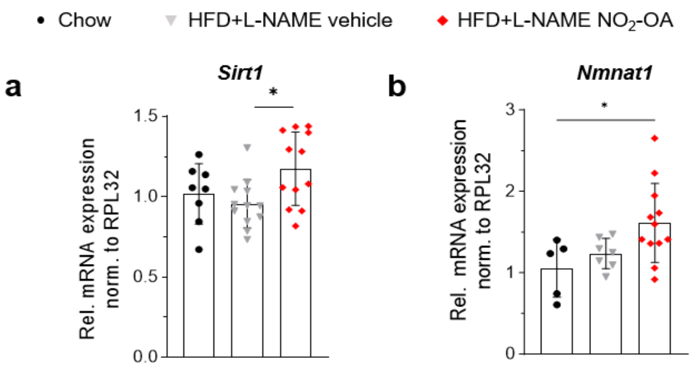


**Figure S6: Downstream targets of AMPK signaling are increased by nitro-oleic acid in HFpEF mice.**
Mice received high fat diet (HFD) and the endothelial nitric oxide synthase inhibitor L-NAME or chow diet (chow) for 15 weeks (wk) and were treated with nitro-oleic acid ($NO_2$-OA) or vehicle for 4 wk. mRNA expression of the NAD-dependent protein deacetylase sirtuin-1 (*Sirt1*) and the nicotinamide nucleotide adenyl transferase 1 (*Nmnat1)* was performed in left ventricular mouse tissue.

### Figure S7

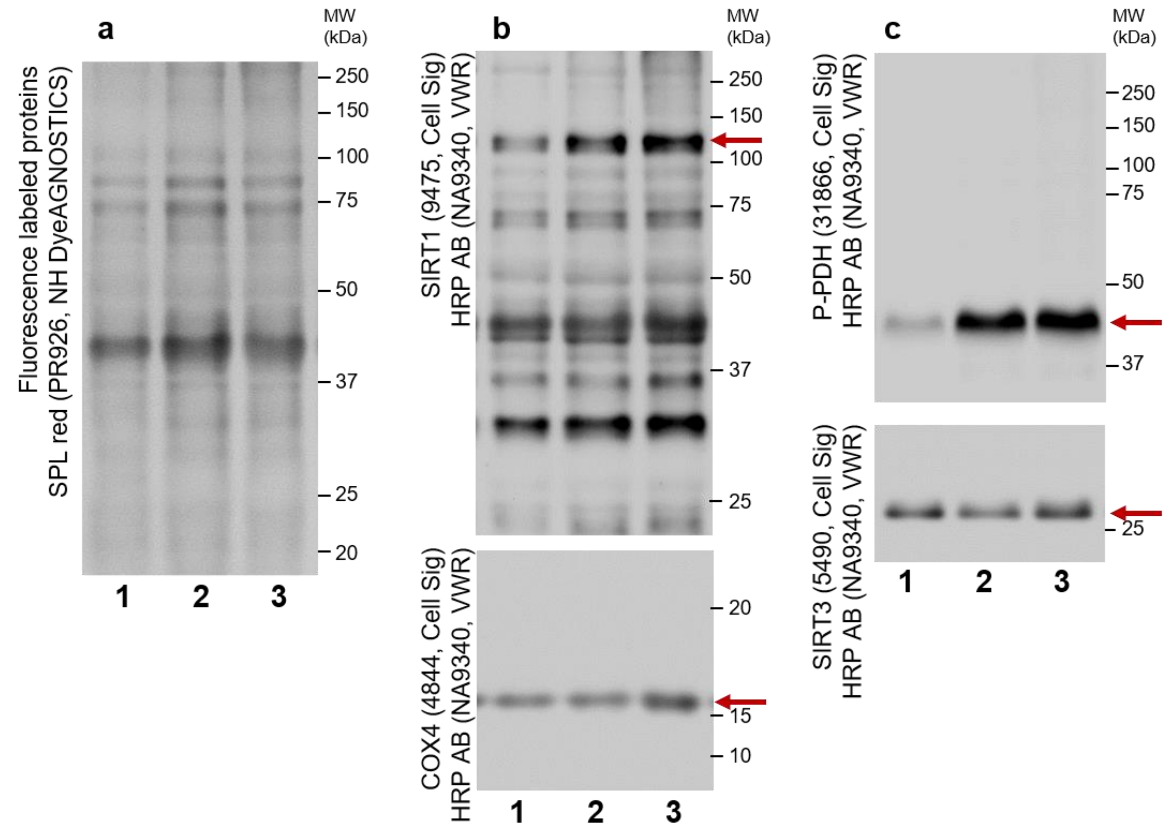


**Figure S7: Detection of protein level in left ventricular tissue of control and HFpEF mice.**
Mice received high fat diet (HFD) and the endothelial nitric oxide synthase inhibitor L-NAME or chow diet (chow) (N=9) for 15 weeks (wk) and were treated with nitro-oleic acid ($NO_2$-OA) (N=6) or vehicle (N=8) for 4 wk (wk 12 to 15). Proteins were isolated and used for proteom analysis and immunoblot. Representative immunoblots are shown for the three experimental groups (1: chow; 2: HFD+L-NAME vehicle; 3: HFD+L-NAME $NO_2$-OA). **a** Proteins were labeled with SPL red to assess total protein amount by fluorescence. **b** The membrane was cut and incubated with sirtuin 1 (SIRT1) (above) or cytochrome c oxidase subunit 4 (COX4) (down) primary antibody. After incubation with HRP-conjugated secondary antibody the protein level was assessed by chemiluminescence. **c** Reblot of membrane and second incubation with antibody detecting the phosphorylation at serine 293 of pyruvate dehydrogenase (P-PDH) (above) or sirtuin 3 (SIRT3) (down). After incubation with HRP-conjugated secondary antibody the protein level was assessed by chemiluminescence.

### Figure S8

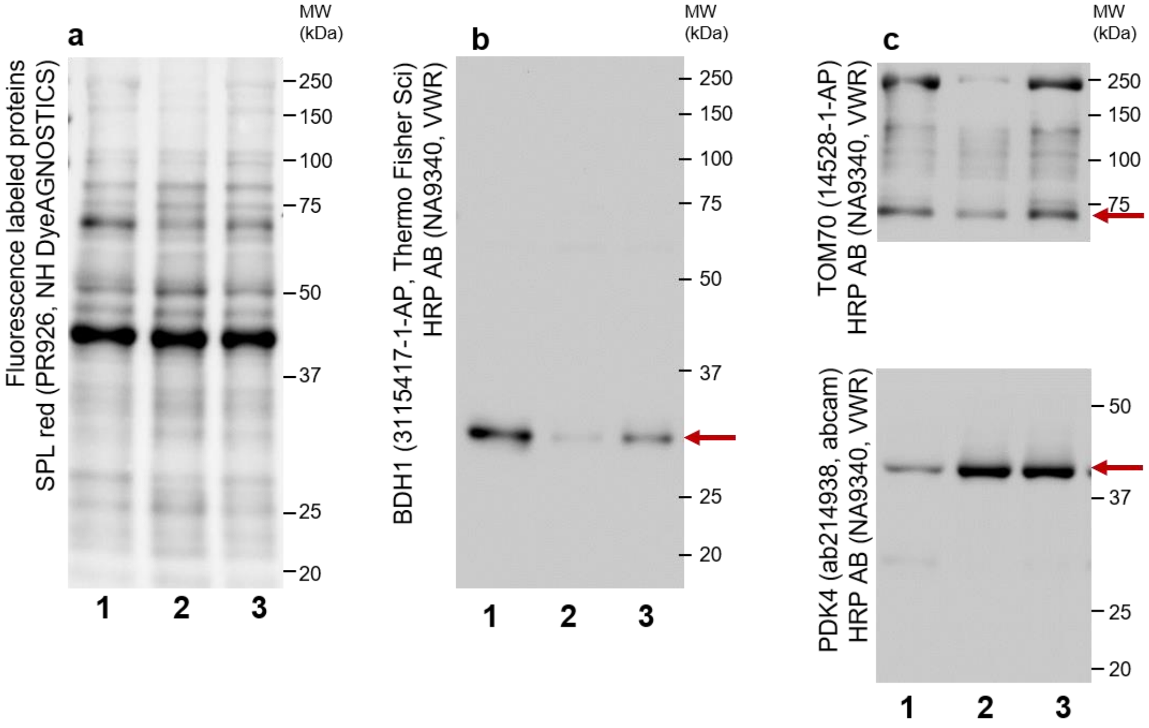


**Figure S8: Detection of protein level in left ventricular tissue of control and HFpEF mice.** Mice received high fat diet (HFD) and the endothelial nitric oxide synthase inhibitor L-NAME or chow diet (chow) (N=9) for 15 weeks (wk) and were treated with nitro-oleic acid ($NO_2$-OA) (N=6) or vehicle (N=8) for 4 wk (wk 12 to 15). Proteins were isolated and used for proteom analysis and immunoblot. Representative immunoblots are shown for the three experimental groups (1: chow; 2: HFD+L-NAME vehicle; 3: HFD+L-NAME $NO_2$-OA). **a** Proteins were labeled with SPL red to assess total protein amount by fluorescence. **b** The membrane was incubated with 3-hydroxybutyrate dehydrogenase 1 (BDH1) primary antibody. After incubation with HRP-conjugated secondary antibody the protein level was assessed by chemiluminescence. *c* The membrane was cut and re-incubated with antibody against translocase of outer mitochondrial membrane 70 (TOM70) (above) or pyruvate dehydrogenase kinase 4 (PDK4) (down). After incubation with HRP-conjugated secondary antibody the protein level was assessed by chemiluminescence.

### Figure S9

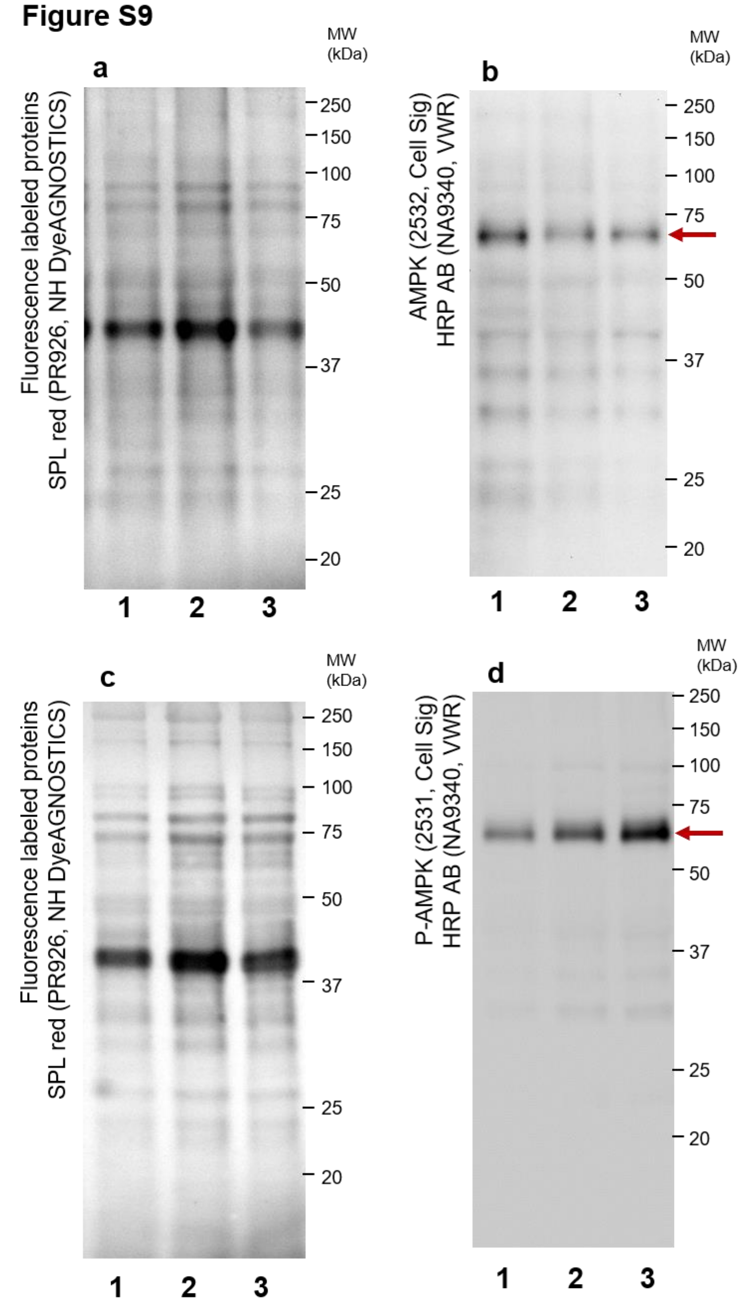


**Figure S9: Detection of protein level in left ventricular tissue of control and HFpEF mice.**
Mice received high fat diet (HFD) and the endothelial nitric oxide synthase inhibitor L-NAME or chow diet (chow) (N=9) for 15 weeks (wk) and were treated with nitro-oleic acid ($NO_2$-OA) (N=6) or vehicle (N=8) for 4 wk (wk 12 to 15). Proteins were isolated and used for proteom analysis and immunoblot. Representative immunoblots are shown for the three experimental groups (1: chow; 2: HFD+L-NAME vehicle; 3: HFD+L-NAME $NO_2$-OA). **a**, **c** Proteins were labeled with SPL red to assess total protein amount by fluorescence. **b** The membrane was incubated with AMP-activated protein kinase (AMPK) primary antibody. After incubation with HRP-conjugated secondary antibody the protein level was assessed by chemiluminescence. **d** The membrane was incubated with antibody detecting the phosphorylation at threonine 172 of AMP-activated protein kinase (AMPK). After incubation with HRP-conjugated secondary antibody the protein level was assessed by chemiluminescence.

**Figure S10**

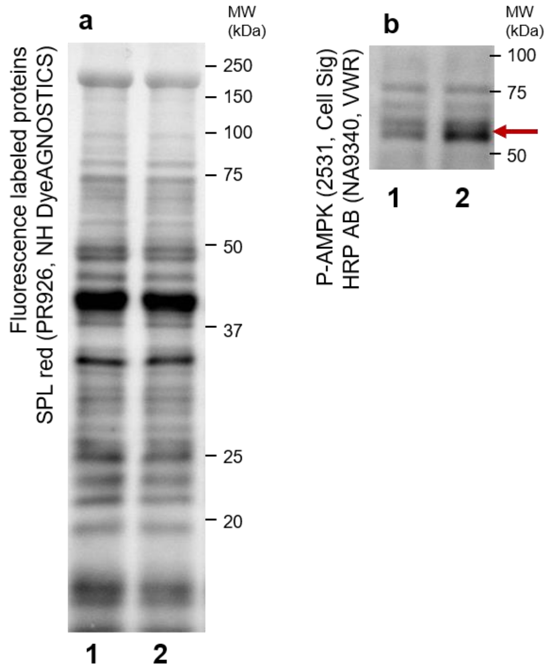


**Figure S10: Detection of protein level in metabolically-stressed cardiomyocytes.**
Isolated adult murine cardiomyocytes were cultivated for 48 hrs under metabolic stress conditions induced by high glucose levels (high Glu), endothelin-1 (ET-1) and hydrocortisone (HC) with treatment of methanol (Vehicle) or nitro-oleic acid ($NO_2$-OA). Representative immunoblots are shown for the two experimental groups (1: high Glu, ET-1, HC vehicle; 2: high Glu, ET-1, HC $NO_2$-OA). **a** Proteins were labeled with SPL red to assess total protein amount by fluorescence. **b** The membrane was cut and incubated with antibody detecting the phosphorylation at threonine 172 of AMP-activated protein kinase (AMPK). After incubation with HRP-conjugated secondary antibody the protein level was assessed by chemiluminescence.
